# MassBank Human: The annotated human metabolome

**DOI:** 10.64898/2026.10.10.758200

**Authors:** Fumio Matsuda, Ryosuke Hayasaka, Taihei Torigoe, Masatomo Takahashi, Yushi Takahashi, Akiyasu C. Yoshizawa, Hiroshi Tsugawa, Shujiro Okuda, Steffen Neumann, Emma L. Schymanski, Yoshihiro Izumi, Akiyoshi Hirayama

## Abstract

Mass spectrometry-based metabolomics is essential for deciphering human metabolic functions and metabolite dark matter. However, the dark matter of the human metabolome is intricately hidden. Therefore, we developed MassBank Human, a curated library that distills 36,512,540 MS2 spectra from 76 public human datasets into 203,886 high-quality spectra via spectral averaging, recalibration, and filtering to achieve high mass accuracy. We further used three metabolite annotation tools and developed a Zipf-Bayesian rank aggregation framework to aggregate their rankings with false discovery rate control. This framework assigned molecular formulas, chemical classes, and planar structures to 55,564, 22,619, and 14,198 spectra, respectively. Approximately half of the annotated structures potentially originated from food, pollutants, or gut microbiota. In a metabolome analysis of colorectal cancer cells, MassBank Human doubled structural annotation coverage (10.6% to 21.2%), identifying acetylated polyamines as markers of metabolic rewiring. This community resource provides an annotated foundation for exploring human metabolomes.

## Introduction

Mass spectrometry-based metabolomics explores the human metabolome to elucidate metabolic function and discover biomarkers^1^. However, this task is complicated because the human metabolome is not a closed system. Various exogenous compounds^2^ and gut microbiota-derived metabolites^3^ link the exposome to endogenous metabolism. Furthermore, the structural annotation of product ion (MS2) spectra is not universal^4^. Therefore, multiple complementary approaches are required to explore this landscape. Spectral matching is constrained by the limited chemical coverage of the reference libraries^5^. Although computational tools generate tentative structural annotations at scale, robust frameworks for controlling the false discovery rate (FDR) remain underdeveloped^6^. Public repositories have accumulated tens of millions of MS2 spectra from human samples. However, the dark matter of the human metabolome is hidden behind extreme scale, redundancy, and intrinsic noise^7^.

As a complementary approach, we revisited and revitalized the concept of the mass spectral tag^8,9^. Mass spectral tags are reproducible, experimentally derived spectra that serve as molecular signatures for the detection of structurally characterized and uncharacterized metabolites^10^. A library of high-quality, human-derived spectra with reliable annotations would constitute an “annotated human metabolome.” Such a library would be an annotation-centric resource that complements MassBank (reference spectrum-centric), Human Metabolome Database (HMDB, metabolite-centric), and Global Natural Product Social Molecular Networking (GNPS, spectrum sharing-centric), which guide both the exploration of metabolite dark matter and annotation of new metabolome data.

In this study, we sought to create a library that distills raw spectra into high-quality consensus spectra based on reprocessed human data-dependent acquisition (DDA) datasets from public repositories—termed the MassBank Human. We further aimed to develop a tool ensemble statistical method that converts consensus across annotation software rankings into FDR-controlled annotation confidence—termed the Zipf–Bayesian rank aggregation (ZBRA) framework. The MassBank Human has the potential to facilitate the annotation of human metabolome data as well as the exposome-aware exploration of the human metabolome. MassBank Human is publicly available via GitHub and a dedicated web portal and will be re-annotated as computational tools evolve.

## Results

### Spectral quality improvement by spectral averaging, recalibration, and filtering

Our reprocessing pipeline accepts human DDA metabolome datasets from public repositories^11-14^. Reprocessing is executed on each DDA dataset individually. The pipeline consists of three sequential modules: preprocessing, annotation, and curation (Fig. 1a). The outputs are a set of MS2 spectra recorded in the standardized MassBank record format (Supplementary Data 1). In addition, mzTab-M files^15^ are generated for each dataset, and all MS2 spectra are stored in MSP files (Fig. 1a). These files are available on GitHub (https://github.com/Shin-MassBank/MassBank-Human) and the MassBank Human database (https://human.massbank.jp/). For example, we obtained 22 data files, including 365,333 MS2 spectra, from Standard Reference Material (SRM) 1950 human plasma using positive-mode DDA on a Q-Exactive instrument. The raw data files were deposited in a data repository (https://repository.massbank.jp/entry/MPST000004). The dataset is denoted as HSA001, its internal identifier in MassBank Human (Supplementary Table 1).

**Figure 1.**
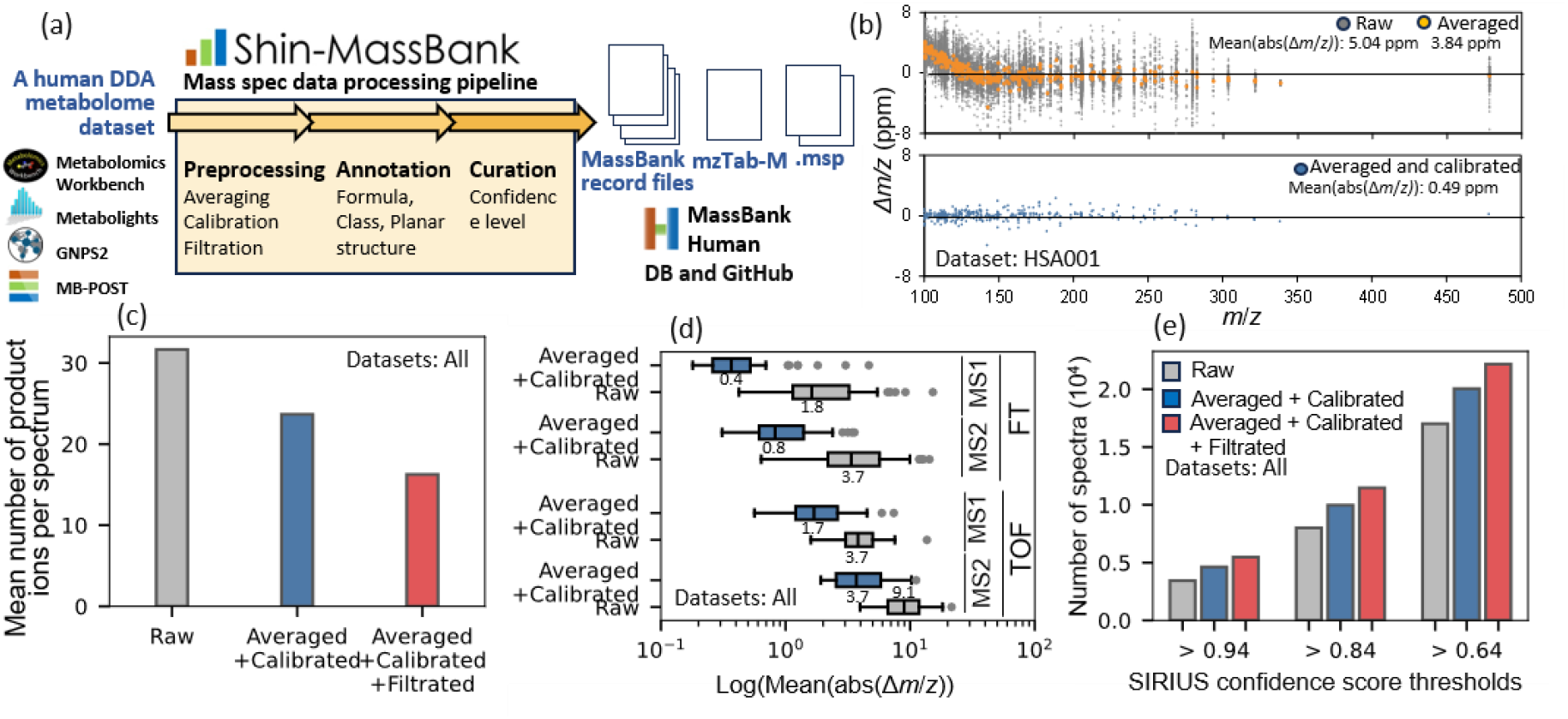
Quality improvement of MS2 spectra by the reprocessing pipeline. (a) The structure of the reprocessing pipeline. (b) Improvement of *m/z* accuracy by spectral averaging and recalibration. The x-axis shows measured *m/z* values in MS2 spectra of the HSA001 dataset; the y-axis shows the deviation from theoretical values (Δ*m/z*). Mean(abs(Δ*m*/*z*)) values are also shown. (c) Reduction in the mean number of product ions per spectrum at each preprocessing tier (means across all datasets). (d) Mass accuracy improvement across datasets. The mean(abs(Δ*m*/*z*)) of MS1 and MS2 spectra was determined for each dataset (Supplementary Table 1). Distributions are shown separately for FT-MS and TOF-MS data; median values are provided in the panel. (e) Improvement of spectral interpretability. All 203,886 spectra were annotated using SIRIUS at each preprocessing tier; bars show the number of spectra exceeding the indicated confidence scores.

DDA datasets are inherently redundant because multiple MS2 spectra are often obtained from the same metabolite^16^. Grouping and averaging these spectra improves the signal-to-noise ratio and *m*/*z* accuracy^17^. For HSA001, averaging 339,576 of the 365,333 spectra (92.9%) yielded 2,873 averaged MS2 spectra. A minimum of 10 spectra per cluster was required for averaging (details in Supplementary Note 1).

We analyzed the mass errors (Δ*m/z*) between experimental and theoretical *m/z* values of product ions to assess the increase in *m/z* accuracy. Querying the 2,873 averaged spectra against the GNPS library^18^ (spectral entropy > 0.7) returned 353 hits, whose formulas were used as pseudo-ground truths because these matches ensured formula annotation with an FDR < 1.0% (Supplementary Note 2). Averaging markedly reduced the dispersion of Δ*m/z* (mean(abs(Δ*m/z*)): 3.84 ppm) relative to raw spectra (5.04 ppm) (Fig. 1b). However, the residual errors showed a systematic hockey stick-shaped drift at low *m/z* region (Fig. 1b), as documented for instrument calibration drift in proteomics and metabolomics^19,20^. We corrected this systematic deviation using piecewise linear calibration curves, thereby substantially improving mass accuracy (mean(abs(Δ*m/z*)): 0.49 ppm) (Fig. 1b). Consequently, the *m/z* matching threshold for database searching was reduced from 8 to 4 ppm for product ions and from 2 to 0.75 ppm for MS1 precursor ions (Extended Data Fig. 1).

We further filtered out false product ions that did not originate from the precursor ions. Because the product ion intensity should correlate with the precursor ion intensity, we calculated the rank correlations between the precursor and product ions across the raw spectra. Product ions that lacked a statistically significant positive correlation with the precursor ion were removed (α = 0.01; Supplementary Note 1).

This pipeline was applied to 76 human DDA datasets. The datasets comprise 34 blood, 19 cells, 14 feces, two heart, two semen, two culture medium, one gingiva, one nasal mucosa and one placenta datasets, indicating a bias toward metabolites found in blood, cells and feces. Of the 36,512,540 initial spectra, 26,336,363 (72.1%) were used to generate 203,886 preprocessed spectra (Supplementary Table 1). Thus, preprocessing reduced the number of spectra to <1% of the original number. Of these, 82.8% were Orbitrap mass spectrometry data (recorded as FT-MS in MassBank records). Each averaged spectrum was assigned a unique identifier. For instance, HSA001P0248000 denotes the average spectrum generated from a spectral cluster with a precursor unit mass of *m/z* 248 in HSA001.

Preprocessing markedly improved spectral quality. The mean number of product ions per spectrum decreased from 31.2 (raw) to 23.4 after averaging and recalibration, and to 16.1 after filtering (Fig. 1c and Supplementary Table 1). This indicates that approximately half of the raw signals represent noise or interference. Mass accuracy improved across all 76 datasets; the mean(abs(Δ*m/z*)) was <0.5 ppm (MS1) and <2.0 ppm (MS2) in most FT-MS datasets (Fig. 1d). To determine whether preprocessing enhanced spectral interpretability, we annotated all 203,886 spectra using SIRIUS in three processing tiers: raw, averaged + calibrated, and fully processed (Supplementary Data 2). The spectrum with the median base-peak intensity in each cluster was used as the representative for the raw tier. High-confidence annotations (score ≥ 0.94) increased from 3,608 (raw) to 4,830 (intermediate) and 5,705 (fully processed), with consistent gains at lower thresholds (Fig. 1e and Supplementary Table 1). This confirms that the preprocessing pipeline significantly enhances spectral interpretability. Thus, preprocessing acts as a distillation because it reduces the vast number of raw spectra while removing noise and contaminants.

### The ZBRA framework: Integrating multi-software rankings under FDR control

All preprocessed spectra were annotated using SIRIUS^21^, MS-FINDER^22^, and MetFrag^23^ to generate candidate rankings for molecular formulae, chemical classes, and planar structures. Herein, "planar structure" denotes the two-dimensional constitution of a molecule without stereochemical configuration, which is identified by the first block of InChiKey^24^. This definition merges stereoisomers (L-leucine and D-leucine, for example) while distinguishing constitutional and positional isomers.

In the query spectrum of HSA001P0248000 (Fig. 2a), 3-hydroxybutyrylcarnitine was ranked first by SIRIUS, MS-FINDER, and MetFrag (1/1/1) (Fig. 2b). A rigorous statistical threshold is required to determine whether such a consensus is sufficient to identify the substance as 3-hydroxybutyrylcarnitine.

**Figure 2.**
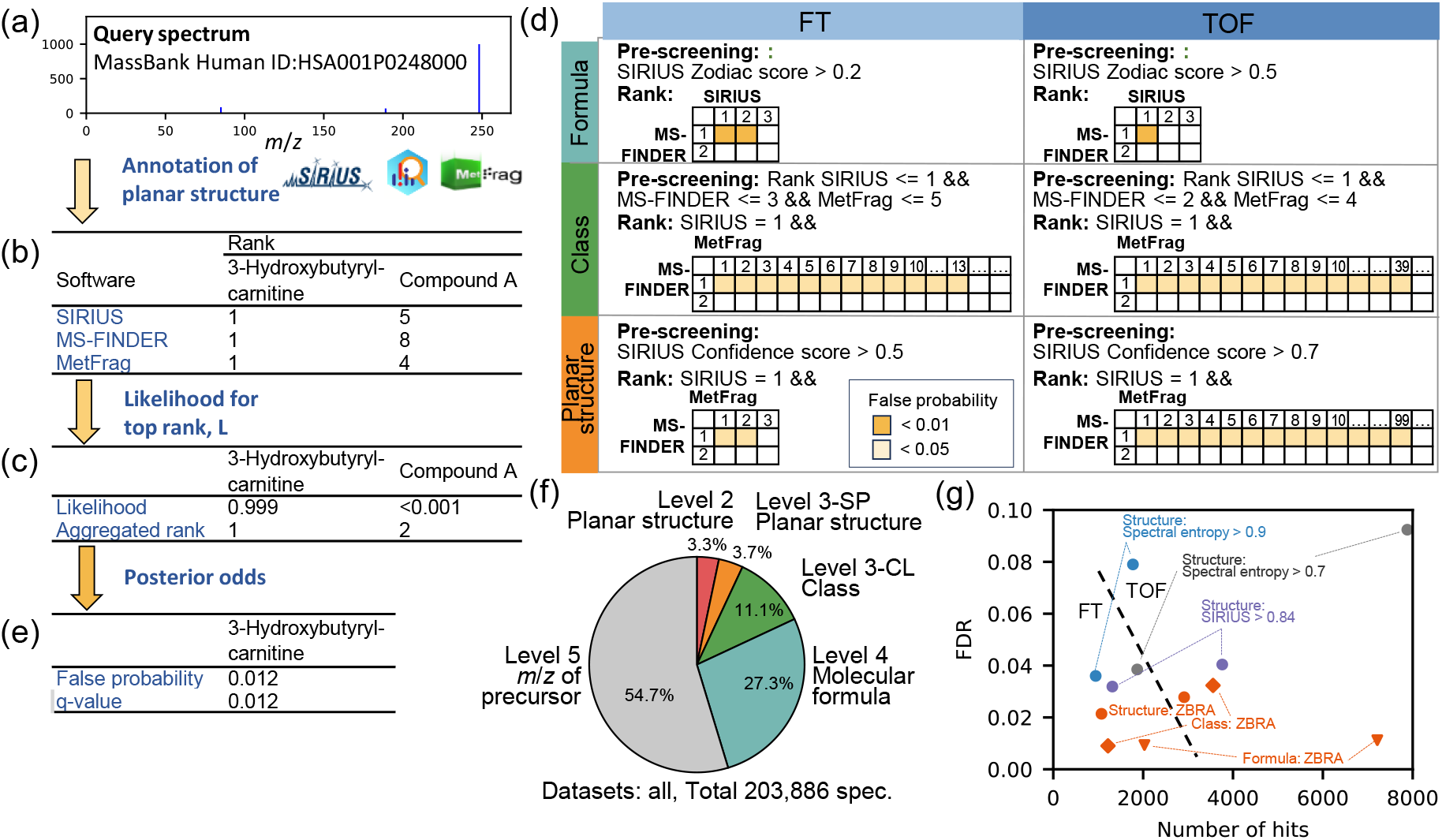
Metabolite annotation using the ZBRA framework. (a) A preprocessed MS2 spectrum, HSA001P0248000. (b) Planar structure rankings from SIRIUS, MS-FINDER, and MetFrag. (c) Rank aggregation. Zipf parameters for each tool were estimated by Bayesian inference; Candidate likelihoods were multiplied across tools and sorted based on their posterior medians to provide the aggregated ranking. (d) Confidence assessment. Shaded regions indicate rank combinations with false probability < 0.01 or < 0.05. Candidates must first pass pre-screening. (e) Confidence evaluation of the annotation; q-values were obtained from false probabilities. (f) Summary of annotation curation. (g) Performance comparison with GNPS spectral-entropy searching and SIRIUS confidence scores. FT-MS and TOF-MS spectra extracted from MassBank were used as evaluation datasets (FT-MS: 3,234 spectra; TOF-MS: 12,464 spectra).

To this end, we developed the ZBRA framework. First, we found that the ranks of the correct hits in the outputs of all three tools followed a Zipf distribution (Supplementary Note 3). The Zipf distribution is an empirical distribution that is observed in various rankings^25^. In Zipf(*k*, *s*) (Eqn (1) in Methods), *k* represents the rank and the parameter *s* indicates the slope of the distribution. A steeper (more negative) *s* indicates better ranking performance.

In this model, Zipf(1, *s*_SIRIUS_) represents the likelihood that the top SIRIUS candidate is a true top hit^25^. The joint likelihood *L* for a candidate ranked first by SIRIUS, MS-FINDER, and MetFrag was calculated as follows:

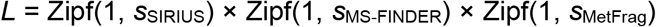

Sorting the candidates using *L* yields an aggregated rank. However, the top-ranked candidate is not guaranteed to be correct, particularly in metabolomics, where the true compound may be absent from the databases used in the annotation software.

Therefore, we modeled the chain of rankings between pairs of tools using a Bayesian approach that estimates the posterior distributions of the Zipf parameters (see Methods). For HSA001P0248000, the aggregated ranking placed 3-hydroxybutyrylcarnitine first (likelihood, 0.999) (Fig. 2c). To determine whether this top hit was correct, we used the posterior distribution of posterior odds (*PO*) computed from the Zipf parameter posteriors. Herein, *PO* is defined as the ratio of the probability that the result is correct to the probability that it is incorrect. The posterior distribution of *PO* for a 1/1/1 ranking by SIRIUS, MS-FINDER, and MetFrag was estimated from the posterior distributions of the Zipf parameters (See Methods). The probability of false annotation (*FP*) was calculated as FP = 1/(1 + *PO*) and the posterior median was used as the representative value.

We mapped the ranking combinations that yielded *FP* < 0.05 for planar structure annotation of the FT-MS data (Fig. 2d). The annotation of HSA001P0248000 was estimated to have *FP* < 0.05 using this map because the spectrum passed pre-screening (SIRIUS confidence score = 0.985; Supplementary Data 1) and the combination 1/1/1 falls within the *FP* < 0.05 region. Within each dataset, the FP values were converted into q-values to control the FDR (Fig. 2e).

Annotations were curated using the annotation confidence level criteria proposed by Schymanski et al.^26^ with modifications. Level 1 (confirmed structure via reference standard matching) could not be applied to ZBRA-based metabolite annotations. Levels 2a (probable structure via exact spectral matching with library databases), 4 (unequivocal molecular formula), and 5 (exact mass of interest) were used without modifications. As the scope of Level 3 (tentative candidate) was too broad, we created sublevels, including Levels 3-SP (tentative annotation by planar structure) and 3-CP (tentative annotation by compound class) (Supplementary Note 4). The annotations were evaluated hierarchically starting from Level 2a. If a planar structure annotation passed the Level 2a threshold (ZBRA q-value < 0.05, spectral entropy > 0.7), the spectrum was assigned to Level 2a. Otherwise, the planar structure (Level 3-SP; ZBRA q-value < 0.05), compound class (Level 3-CP; ZBRA q-value < 0.05), and molecular formula (Level 4; ZBRA q-value < 0.01) annotations were sequentially tested. The remaining spectra were assigned to Level 5. Of the 203,886 averaged spectra, 6,720 (3.3%) were annotated as Level 2a, 7,478 (3.7%) as Level 3-SP, 22,619 (11.1%) as Level 3-CP, and 55,564 (27.3%) as Level 4 (Supplementary Table 2, Fig. 2f). In total, 111,505 spectra (54.7%) remained at Level 5. In this pipeline, HSA001P0248000 (Fig. 2a) was annotated as 3-hydroxybutyrylcarnitine at Level 3-SP. Importantly, Level 3-SP is a tentative annotation that does not rule out other possible planar structures. Thus, it should be interpreted as the most likely isomer among the possible positional and structural isomers (Supplementary Note 4).

We benchmarked ZBRA against existing scoring approaches using annotated MassBank spectra as ground-truth queries (FT-MS: 3,234 spectra; TOF-MS: 12,464 spectra) (Fig. 2g). For planar structures in the FT dataset, ZBRA (q-value < 0.05) yielded 1,073 hits with an empirical FDR of 0.021. In comparison, GNPS spectral searching at spectral entropy thresholds of 0.7 and 0.9 yielded FDRs of 0.039 and 0.036, respectively, and a SIRIUS confidence score threshold of 0.84 yielded a comparable number of hits (1,315) at an FDR of 0.032. Similar trends were observed for the TOF dataset (Fig. 2g). For the molecular formulas (q-value < 0.01) and chemical classes (q-value < 0.05), the empirical FDRs were both 0.009 in the FT dataset. These values were 0.011 and 0.032, respectively, in the TOF dataset. Similar FDR levels were obtained using an independent approach (Supplementary Note 5).

### Construction of MassBank Human

The full pipeline was applied to 76 datasets (5,543 files, see Methods), resulting in a library of 203,886 records (Supplementary Table 3). Each entry stores the preprocessed spectrum, multilevel annotations with confidence criteria, experimental conditions, and biological metadata in the MassBank record format (Supplementary Data 1). Two MSP files containing all the positive- and negative-ion mode MS2 spectra were generated to be used in metabolome data analysis tools. In addition, mzTab-M files were produced for each dataset to describe the precursor ion signal intensities and relationships between the averaged and raw spectra^15^. We designated this resource as MassBank Human and deposited all files on GitHub (https://github.com/Shin-MassBank/MassBank-Human) and a dedicated web portal (https://human.massbank.jp/) (Fig. 1a).

### Application 1: Exploration of the human metabolome

The 14,198 Level 2a and 3-SP spectra encompassed 2,622 unique planar structures (Supplementary Table 3). The ChemOnt ontology used in ClassyFire^27^ categorized 1,179 planar structures (44.3%) as amino acids, peptides, and analogs, followed by lipids and lipid-like molecules (610 planar structures, 22.9%) (Fig. 3a, Supplementary Table 3). This distribution partly reflected the predominance of large fecal and blood (serum/plasma) datasets in the library. Next, we traced metabolites potentially derived from exogenous sources. Planar structures derived from the food, gut microbiota, and environmental pollutants were identified by searching FooDB^28^, MiMeDB^29^, and DrugMedicInfo in PubChemLite^30^, respectively. Moreover, filtering SMILES containing an ethylenedioxy (OCCO) moiety yielded 122 polyethylene glycol derivatives. We identified1,078 food-derived metabolites, 602 gut microbiota-derived metabolites, and 672 environmental pollutants (Fig. 3a). However, it is important to note that the DrugMedicInfo category contains an amount of bioactive ingredients derived from biological sources. Although these are rough estimates, approximately half of the detected planar structures potentially originated from sources other than endogenous human metabolism^31^.

**Figure 3.**
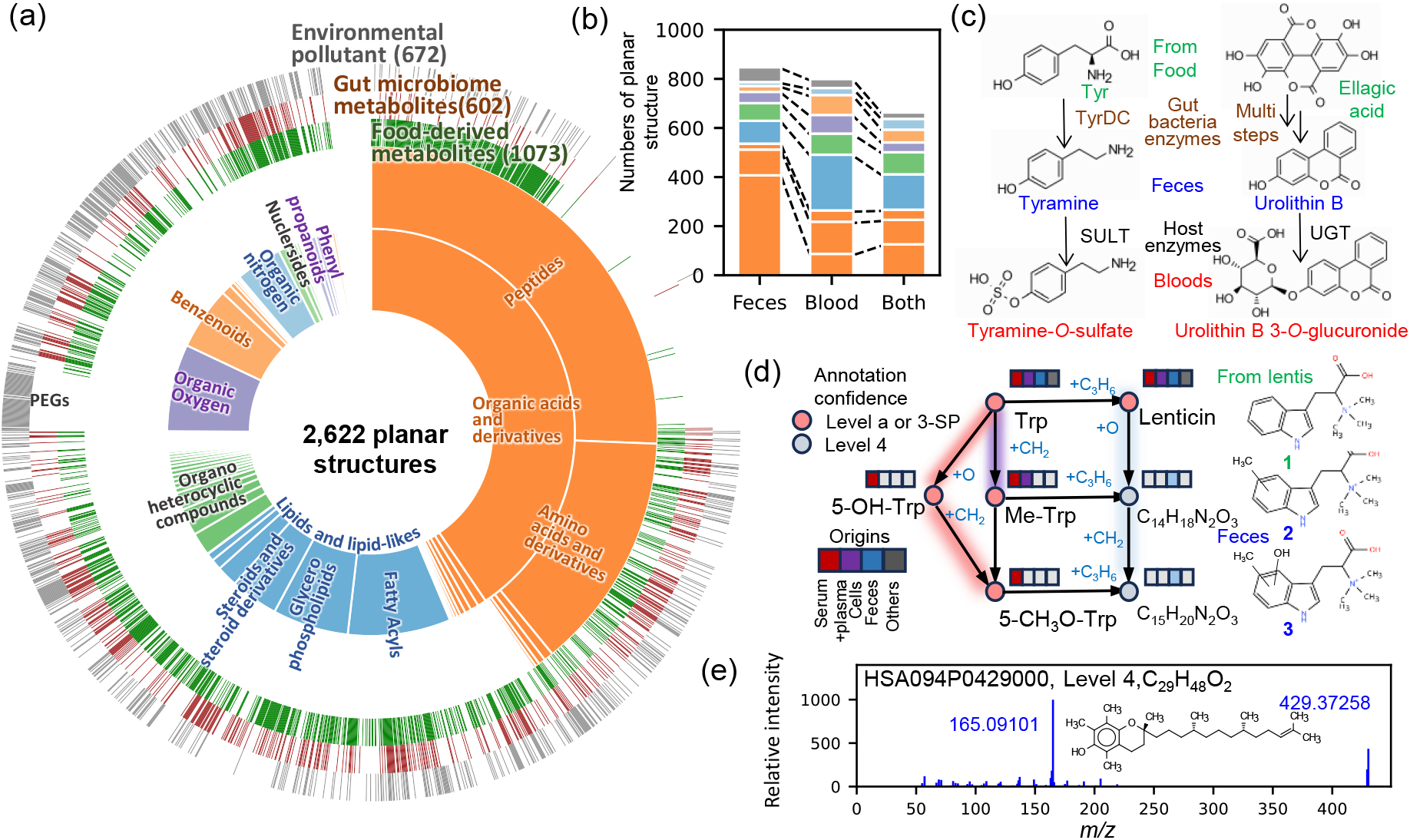
Exploration of the human metabolome using MassBank Human. (a) Chemical-class composition of the 2,622 unique planar structures (14,198 Level 2a or 3-SP spectra), as determined using ChemOnt. Overlaid bars indicate structures attributed to food (green), the gut microbiome (brown), and environmental pollutants (gray) identified using FooDB, MiMeDB, and PubChemLite (DrugMedicInfo), respectively. (b) Planar structures detected exclusively in feces, blood, or in both; colors indicate the chemical classes defined in (a). (c) The interplay between food-derived metabolites, the gut microbiome, and host metabolism. Metabolites detected exclusively in feces or blood are shown in blue and red, respectively; putative food-derived precursors are shown in green; estimated metabolic pathways are indicated. TyrDC, tyrosine decarboxylase; SULT, sulfotransferase; UGT, UDP-glucuronosyltransferase. (d) The molecular subnetwork around tryptophan and lenticin. Each node represents a cluster of highly similar spectra (cosine product score > 0.7) sharing an identical precursor; node color indicates the best annotation confidence level in the cluster, and node labels list the sample types (blood, cell culture, and feces) in which the spectra were detected. Edges connect nodes containing at least one spectral pair with a modified cosine product score > 0.7. Structures of lenticin (**1**) and its deduced methylated (**2**, C_15_H_20_N_2_O_2_) and hydroxylated (**3**, C_15_H_20_N_2_O_3_) derivatives are shown. (e) MS2 spectrum of one of the most-wanted unidentified spectra, recurrently detected in 20 of the 76 datasets. The spectrum was tentatively annotated as α-tocomonoenol through manual curation of the molecular network (Supplementary Note 6).

Matrix-specific analysis identified 874 structures related to feces and 821 in blood (Fig. 3b); peptides and lipids were predominant in the feces (407 peptides) and blood (227 lipids), respectively. These structures allowed us to directly probe the interplay between gut microbial and host metabolism. Gut bacteria produce metabolites such as indole, which are absorbed and converted into indoxyl sulfate by host enzymes^32^. Therefore, we searched for metabolites detected exclusively in feces whose conjugates were detected exclusively in the blood. We have identified two precursor–conjugate pairs: tyramine and tyramine *O*-sulfate, and urolithin B and urolithin B 3-*O*-glucuronide (Fig. 3c). Tyramine is formed from tyrosine residues by host and bacterial tyrosine decarboxylases in the gut^33^. The restriction of tyramine to the feces, together with the restriction of its sulfate to the blood, suggests the absorption of microbial tyramine, followed by host sulfotransferase-mediated conjugation. Urolithin B is produced from dietary ellagitannins and ellagic acid via a multistep gut bacterial transformation^34^. Its distribution showed a corresponding pattern, suggesting absorption and host UDP-glucuronosyltransferase-mediated glucuronidation. Thus, the human metabolome emerges from the interplay between dietary inputs, gut microbial transformation, and host conjugative metabolism^35^.

To further investigate this interplay, we constructed an inter-dataset molecular network^36^ including Level 2a, 3-SP, 3-CP, and 4 spectra (Extended Data Fig. 2). We subsequently focused on the subnetwork surrounding tryptophan and lenticin (**1**). Lenticin, a trimethylated tryptophan (Fig. 3d), is not synthesized by humans. Therefore, it is used as a biomarker of lentil intake^37^. The lenticin node was directly connected to tryptophan, and two sequentially connected Level 4 nodes (C_15_H_20_N_2_O_2_ and C_15_H_20_N_2_O_3_) were assigned to its methylated (**2**) and hydroxylated (**3**) derivatives by MS2 inspection (Extended Data Fig. 3a). Neither structure has been reported as a human metabolite. Biological metadata revealed that lenticin was detected in diverse datasets (blood, cell culture, and feces) (Fig. 3d, Extended Data Fig. 3b). However, it was absent in the two Chinese cohort datasets, likely reflecting dietary differences among countries. Furthermore, tryptophan analogs were abundant in blood and cell cultures, whereas lenticin derivatives were restricted to a US fecal cohort (HSA010)^38^. In this dataset, the MS2 spectra of lenticin were obtained from 18 of the 92 samples. The high inter-individual variation in the profiles of lenticin-derived analogs suggests gut microbiota-mediated transformation (Extended Data Fig. 3c).

Finally, MassBank Human highlighted issues that remain to be solved. Projection of HMDB^39^ and MassBank Human structures onto the MS-FINDER compound space (554,833 compounds) by t-SNE using MACCS keys revealed dense clustering in localized regions (Extended Data Fig. 4). This reflected both the structural bias of human metabolism and hard-to-annotate metabolite classes. Beyond mapping this bias, the library prioritizes recurrent unidentified spectra that are detected across many datasets and are consequently high-priority targets for structure elucidation (Supplementary Table 4). For example, the spectrum in Fig. 3e was detected in 20 of the 76 datasets but remains at Level 4 because SIRIUS, MS-FINDER, and MetFrag failed to reach a consensus, ranking a vitamin E analog, cholesteryl acetate, and α-tocomonoenol at the top, respectively. Analysis of the subnetwork tentatively identified the compound as α-tocomonoenol, an α-tocopherol analog bearing one additional side-chain double bond (Supplementary Note 6). Moreover, fragmentation analysis assigned planar structures of α-tocopherol derivatives to 13 additional spectra. Thus, MassBank Human enables the progressive exploration of the uncharted human metabolome.

### Application 2: Expanding annotation coverage in metabolome data

MassBank Human can serve as a supplementary spectral library for the annotation of human metabolome data, enabling the use of both structurally identified and unidentified spectra previously detected in humans, together with their biological context.

We evaluated this capability through non-targeted metabolome analysis of cultured cancer cells. A DDA dataset was newly acquired from HCT116 human colorectal cancer cells and an *IDH1*-mutant line (R132H/+) using HILIC-MS/MS to model oncogenic metabolic rewiring^40^. Processing with MS-DIAL yielded a data matrix of 2,412 features (Supplementary Data 3). Standard MS-DIAL spectral library searching (324,191 reference spectra) annotated 254 features (10.6%) above the score threshold, whereas 1,105 features (45.9%) had low scores and 43.5% remained unscored (Fig. 4a). Supplementing MS-DIAL with the MassBank Human library (MSP format) annotated an additional 45 (1.9%), 211 (8.7%), 87 (3.6%),177 (7.3%), and 256 (10.6%) features at Levels 2a, 3-SP, 3-CP, 4, and 5, respectively (Fig. 4b). This doubled the total number of structural annotations.

**Figure 4.**
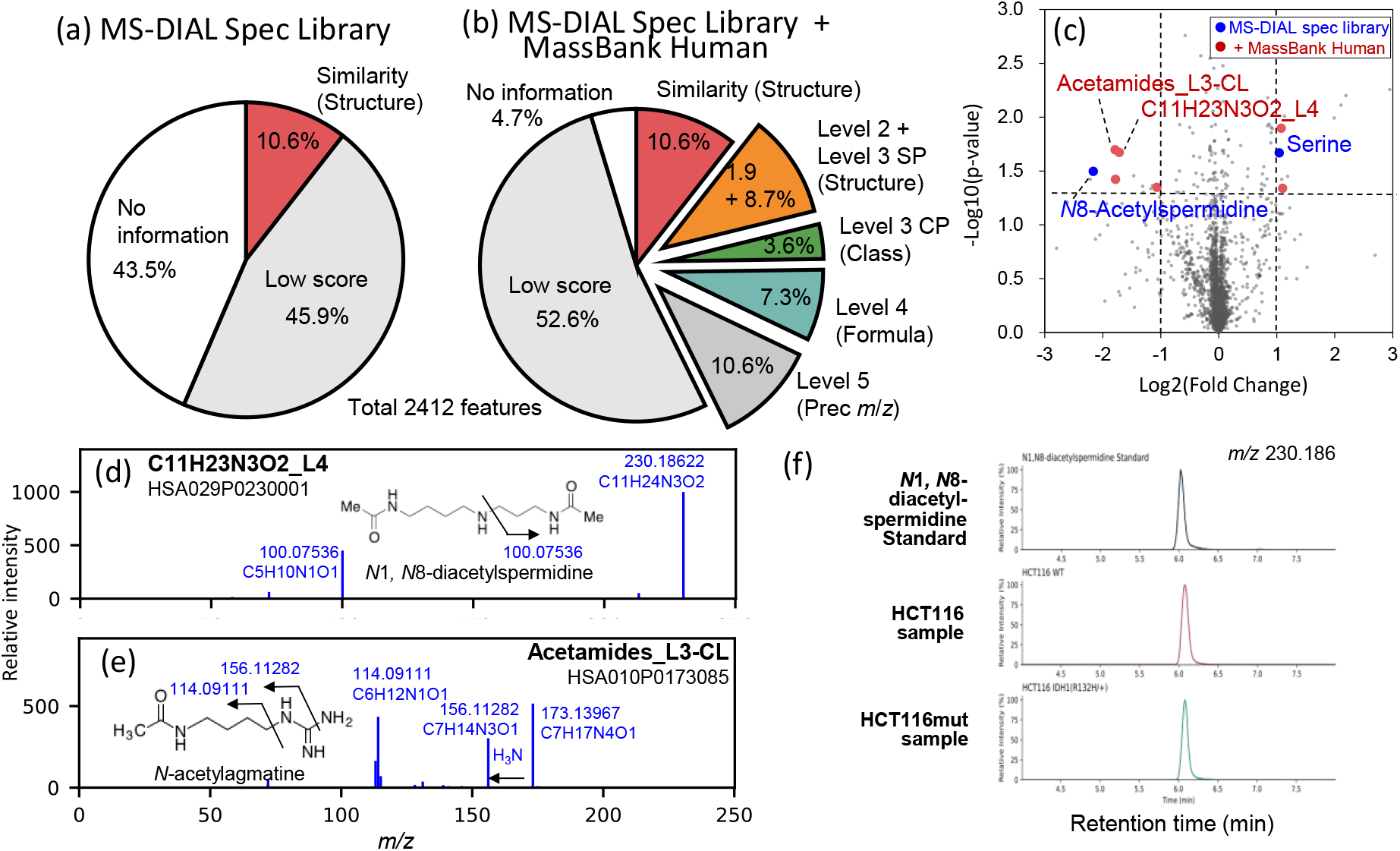
Expansion of annotation coverage in colorectal cancer cell metabolomics using MassBank Human. A metabolome dataset was obtained from parental and *IDH1* (R132H/+) HCT116 cells; MS-DIAL processing produced 2,412 features. (a) Annotation by the MS-DIAL spectral library alone; segment size indicates the fraction in each criterion (above the default similarity threshold, low score, unannotated). (b) Annotation after adding MassBank Human; new segments show features gained at Levels 2a, 3-SP, 3-CP, 4, and 5. (c) A volcano plot comparing parental and *IDH1* (R132H/+) cells; blue and red dots represent significantly altered features annotated by the MS-DIAL library and MassBank Human, respectively. (d, e) MS2 spectra of the C_11_H_23_N_3_O_2_ (Level 4) and acetamide (Level 3-CP) features from (c), with estimated product ion assignments for (d) *N*1,*N*8-diacetylspermidine and (e) *N*-acetylagmatine. (f) Overlaid extracted ion chromatograms of m/z 230.186 for the authentic standard and cell samples (parental and mutant); the identical retention time (∼6.08 min) confirms chromatographic identity.

Differential analysis identified 21 altered features in the IDH1-mutant cells (Supplementary Table 5, Fig. 4c). Standard MS-DIAL annotated two of these features, whereas MassBank Human assigned the matching spectra above the threshold to six additional features. Among these, one spectrum (HSA029P0230001) was curated at Level 4 (C_11_H_23_N_3_O_2_) using this pipeline (Fig. 4d). Inspection of its MassBank record led us to annotate this spectrum as *N*1,*N*8-diacetylspermidine based on the top-ranked SIRIUS candidate, interpretation of the MS2 spectrum (Fig. 4d), and biological context (detection in the human cancer cell dataset HSA029)(Supplementary Data 4). This annotation was supported by the reported status of this polyamine as a urinary tumor marker for breast and colorectal cancers^41^. Similarly, a feature annotated as acetamide at Level 3-CP was assigned to *N*-acetylagmatine (Fig. 4e). An authentic standard confirmed identical retention times and MS/MS spectra for *N*1,*N*8-diacetylspermidine (Fig. 4f, Extended Data Fig. 5). These findings underscore the ability of MassBank Human to expand annotation coverage and to convert it into biological insights.

## Discussion

In this study, we demonstrate that MassBank Human, a spectral library generated from multiple human metabolome DDA datasets, systematically identified dark matter metabolites (Fig. 3) and expanded the annotation coverage in human metabolomics (Fig. 4). By exploiting the spectral redundancy inherent to DDA, our reprocessing pipeline distilled 36,512,540 raw MS2 spectra from 76 public datasets into 203,886 high-quality averaged spectra through spectral quality improvement (Fig. 1) and statistical FDR control (Fig. 2). To our knowledge, the resulting library is the largest quality-controlled and systematically annotated MS2 resource dedicated to the human metabolome; it preserves repository-derived spectra and their biological context. MassBank Human complements compound-centric knowledge bases such as HMDB^39^ and community spectral repositories such as MassBank^42^ and GNPS^18^.

In practical applications, MassBank Human doubled the structural annotation coverage (10.6%–21.2%) for cancer cell profiling (Fig. 4a,b). The structural annotations and contextual clues recorded in each spectrum indicated that acetylated polyamines are markers of IDH1-driven metabolic rewiring (Fig. 4d,e). For instance, the annotation of *N*1,*N*8-diacetylspermidine is consistent with the reported status of this polyamine as a urinary tumor marker for breast and colorectal cancers^41^. Spermidine and agmatine are polyamines synthesized from arginine and ornithine, respectively. Their supply is coupled to glutamate and, hence, to 2-oxoglutarate metabolism^43^. Therefore, depletion of their acetylated forms is consistent with the rewiring of 2-oxoglutarate metabolism caused by the *IDH1* mutation. The present findings demonstrate that unidentified spectra accumulated across studies can be converted into biological discoveries.

Moreover, the meta-analysis of the integrated multi-study datasets in MassBank Human enhanced the resolution of the molecular network and enabled biological context-driven interpretation, as demonstrated for tryptophan and tocopherol derivatives (Fig. 3d and Supplementary Note 6). The annotated landscape indicates that approximately half of the detected planar structures potentially originated from food, environmental exposure, or the gut microbiota (Fig. 3a), indicating that the human metabolome must be interpreted as a superposition of endogenous metabolism and exposome^3,31^.

The ZBRA framework enables the FDR control of molecular formulas, chemical classes, and planar structures (Fig. 2). Rank aggregation and error rate estimation have precedents in computational metabolomics. A consensus among multiple in silico tools improves candidate ranking^44,45^. Statistical rank-aggregation methods have been developed for noisy ranked lists in genomics^46^. FDR estimation has been reported for formula search, spectral library matching^6^, formula annotation^47^, and single tool structure annotation^48^, paralleling multi-engine integration in proteomics^49^. ZBRA unifies these separate strands into a single framework with three distinctive features. First, it explicitly models the rank distribution of correct hits with the Zipf likelihood rather than a uniform null (Supplementary Note 3). Second, it derives the posterior distributions of the likelihood ratio from the Bayesian inference of rank chains between tools, consequently requiring no decoy construction (Fig. 2). Third, it applies a consistent FDR-controlled criterion across molecular formulas, chemical classes, and planar structures (Fig. 2). Its tool ensemble design allows confidence estimates to improve as ranking-based annotation software evolves. Conversely, parameter estimation requires substantial data for calibration, making the framework more suitable for repository-scale reanalysis than for isolated single-study use.

The constructed library has several limitations. Large blood, cell, and fecal cohorts dominate the sample types. Therefore, chemical class distributions (Fig. 3a) should not be read as unbiased estimates of the human metabolome. Spectral averaging with the minimum cluster size (*n* = 10) inevitably discards rare low-frequency spectra. Moreover, more than half (54.7%) of MassBank Human spectra currently lack molecular formula assignments (Fig. 2f). These constraints will be partly relaxed as the repository content grows in scale and diversity, and as the nature of hard-to-annotate spectra is analyzed further such as considering in-source fragmentation^50^.

MassBank Human could not have been constructed without public metabolome repositories, deposited datasets, annotation software packages, and standardized formats established by MassBank and HUPO-PSI. The Shin-MassBank project will regularly re-annotate repository datasets using emerging software tools to maintain current resources, progressively converting unannotated spectra into annotated entries under the same FDR-controlled framework. Continued community-driven data sharing, standardized reporting, and automated reprocessing will be imperative for fully resolving the dark matter of the human metabolome.

## Methods

### Collection of human DDA metabolome datasets

The data files used in this study were retrieved from public metabolomic data repositories, including Metabolomics Workbench (https://www.metabolomicsworkbench.org/)^11^, MetaboLights (https://www.ebi.ac.uk/metabolights/)^12^, GNPS2 (https://gnps2.org/homepage)^13^, and MB-POST (https://repository.massbank.jp/)^14^. Datasets acquired from human samples using the DDA were downloaded based on their metadata. Each MassBank Human dataset was assigned a unique ID (e.g., HSA001) and consisted of DDA data files acquired using a single analytical method in a single polarity (Supplementary Table 1). In total, 76 datasets comprising 5,543 data files were used (TOF-MS: 22 datasets and 1,126 files; Orbitrap (FT)-MS: 54 datasets and 4,417 files).

### Software and resources for the reprocessing pipeline

The reprocessing pipeline was entirely implemented in Python 3.12. The HiveSDK_0.19.0_JA library provided by Reifycs Inc. (https://www.reifycs.com/products/hive/) was used to access data files in Hive format. Additionally, MetFrag 2.6.1^23^, MS-FINDER 3.70^22^, and SIRIUS 6^21,51^ were used for compound annotation. The parameters used are summarized in Supplementary Data 6. For SIRIUS 6, default settings were used for compound database selection. MsfinderStructureDB-VS15. esd supplied by MS-FINDER and supplemented with a custom database that fills gaps relative to SIRIUS 6 was used for MS-FINDER and MetFrag (Supplementary Data 7). For MetFrag, the PubChemLite database (version 1.41.0, https://doi.org/10.5281/zenodo.5995885) was used as another compound database^52^.

MassBank record data were obtained from the MassBank GitHub repository (retrieved December 10, 2024, https://doi.org/10.5281/zenodo.14221628)^42,53^. The GNPS spectral library (ALL_GNPS_NO_PROPOGATED.json, retrieved on November 6, 2023) was downloaded from https://external.gnps2.org/gnpslibrary^13^.

Conversion from the InChIKeys to ChemOnt categories was performed using InChIKeyClassyfireDB-VS5.Icd, which was included with MS-FINDER and the ClassyFire API (https://bitbucket.org/wishartlab/classyfire_api/src/master/)^27^. The ChemOnt definition file (ChemOnt_2_1.obo) was obtained from the official ClassyFire website (http://classyfire.wishartlab.com/downloads).

For similarity searches against the GNPS spectral library, candidate entries matching the precursor ion *m*/*z* within a specified tolerance were extracted (MS1 threshold: 10 ppm for TOF and 3 ppm for FT), followed by evaluation of MS2 spectral similarity using the spectral entropy score (MS2 threshold: 0.02 Da)^54^.

t-SNE was performed using sklearn.manifold.TSNE in the scikit-learn package (https://scikit-learn.org/stable/). MACCS fingerprints were obtained using RDKit (version 2026.03.1; https://www.rdkit.org). The molecular network was constructed by modifying the original method^55^. For each pair of Level 2a, 3-SP, 3-CP, and 4 spectra, spectral similarity (modified cosine score > 0.7) was evaluated by direct comparison of the product ion formulas. The formula differences between C1H2, C2H4, H2, O1, H2O1, C3H6, and C2H2O1 were used to generate a simplified network. The molecular network was visualized using Cytoscape 3.10.3^56^.

### Data reprocessing pipeline

Reprocessing was conducted for each dataset (Expanded Data Fig. 6). The reprocessing pipeline consisted of four major steps: averaging (Step 1), recalibration and filtering (Step 2), annotation (Step 3), and curation (Step 4). First, a RecordTemplate.txt file was created for each dataset to describe the metadata, threshold values used for averaging, and calibration curves (Supplementary Data 5). The output of the reprocessing pipeline is a set of MassBank record files. Each MassBank record file has a unique ID and contains only one processed spectrum. Annotation- and curation-related information was recorded in an extended format using the COMMENT field of the MassBank record format (Supplementary Table 6).

### Step 1: Averaging

Raw data files from each dataset were converted to the Hive format before spectral averaging. Spectral averaging was performed as previously described^57^, with modifications. All MS2 spectra from all the data files were gathered into a spectrum pool. A seed spectrum was selected in descending order of base-peak intensity, and all spectra satisfying |Δretention time| < 1.5 min, precursor ion |Δ*m/z*| < ms1_mz_raw_thres = 1.5 ppm, and cosine product score > 0.9 were collected. A threshold of ms2_mz_raw_thres = 8 ppm was used for the spectral comparison. These thresholds were configured manually for each dataset. When ≥10 similar spectra were collected, the intensities and *m/z* values were averaged across the clusters of the raw spectra. Product ions observed in <70% of the raw spectra in a cluster were discarded^57^. In addition, spectra used for averaging were removed from the spectrum pool. This averaging procedure was iterated in descending order of the base-peak intensity until no candidate seed spectra remained. The same averaging process was applied to the MS1 spectra corresponding to each product ion spectrum.

### Step 2: Recalibration and filtering

In the first iteration of *m/z* recalibration, similarity searches against the GNPS spectral library were performed using uncalibrated averaged spectra as queries. Top-scoring hits with scores >0.7 were considered positive matches. The formulae of these hits were used as pseudo-ground truths because such matches ensure formula annotation with an FDR < 1.0% (Supplementary Note 2). Using the molecular formula of the hit compound as the upper bound, molecular formulas were assigned to the product ions by selecting the formula closest to the observed *m/z* value. Mass errors (Δ*m/z*), defined as the difference between theoretical and observed *m/z* values, were calculated and plotted against *m/z* (*m/z* on the x-axis, Δ*m/z* on the y-axis). Calibration curves for *m/z* correction were constructed from these plots using linear interpolation (ms1_mz_correction and ms2_mz_correction; Supplementary Data 5). In the second iteration, *m/z* values were corrected using the generated calibration curves; error plots were generated in the same manner. Tolerance thresholds for molecular formula searching during annotation were set based on the dispersion of Δ*m/z* in the corrected plots. An identical procedure was performed for the MS1 spectra. The determined thresholds were recorded in RecordTemplate.txt as ms1_mz_averaged_thres and ms2_mz_averaged_thres and used in the subsequent steps.

For each product ion, the Spearman rank correlation coefficient between the intensities in the raw spectra and corresponding precursor ion intensities was calculated. Product ions without a significant positive correlation were removed using a one-sided test (null hypothesis: correlation coefficient = 0, α = 0.01). Unique IDs were assigned to the preprocessed spectra, which were saved in the MassBank record format^42^. For example, "HSA001P0205016" indicates the averaged product ion spectrum with a precursor ion unit mass of m/z 205 in positive mode (P) of the HSA001 dataset, derived from the 16th largest cluster of similar spectra used for averaging.

### Step 3: Annotation

Annotation was performed using MetFrag, MS-FINDER, and SIRIUS with the preprocessed MS1 and MS2 spectra as queries. The mass tolerance thresholds recorded in ms1_mz_averaged_thres and ms2_mz_averaged_thres were applied. Planar structural identity was evaluated by matching the first block of the InChIKey. Chemical class annotations with ChemOnt categories were retrieved via InChIKeys using ClassyFire. When duplicate chemical classes appeared in the ranking results, non-top-ranked duplicate entries were removed before reranking. In addition to the software-based annotation, a spectral similarity search was performed against the GNPS spectral library using the spectral entropy score. The annotation results from each software tool were extracted and recorded in the COMMENT: PROPOSED_MF, COMMENT: PROPOSED_COMPOUND, and COMMENT: PROPOSED_CLASS fields of each MassBank record file.

### Preparation for Step 4: Bayesian estimation of the rank-chain model parameters

After completing Step 3 for all 76 datasets, the combined annotation results were used to prepare for Step 4. We assumed that the ranks of the correct hits in the outputs of SIRIUS, MS-FINDER, and MetFrag follow a Zipf distribution (validated in Supplementary Note 3). The Zipf distribution (Eqn (1)) is an empirical distribution observed across diverse ranking systems^25^:

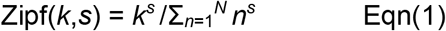

where *k*, *s* (< −1), and *N* represent the rank, exponent governing the slope, and total number of entries in the ranking, respectively; the denominator normalizes the probabilities to sum to 1. *N* = 200 was used throughout this study.

Zipf(*k*, *s*) can be interpreted as the likelihood of a candidate of rank *k* being the top hit^25^. When a candidate is ranked *k*, *l*, and *m* using SIRIUS, MS-FINDER, and MetFrag, respectively, the aggregated likelihood *L* is determined as follows:

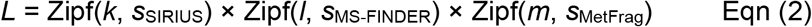

Sorting the candidates using *L* yields an aggregated ranking. However, the top aggregated candidate is not guaranteed to be correct in non-targeted metabolomics because the correct compound is not necessarily contained in the searched compound database^4,48^.

We modeled the chain of rankings between the pairs of tools to determine whether a top-ranked candidate was correct. An example is a chain of rankings between a planar structure ranked first by SIRIUS and its rank assigned by MS-FINDER. Because a planar structure ranked first by SIRIUS tends to also rank highly in MS-FINDER, the rank distribution observed across a dataset resembles "All" in Extended Data Fig. 7. A top-ranked SIRIUS hit belongs to one of three scenarios: true hit (T), false hit (F), or true planar structure is out of the database (O) (Extended Data Fig. 7). We denoted the mixture proportion of query spectra whose true compound is present in the candidate ranking by *π*_DB_; *π*_DB_ is not fixed a priori but is treated as a model parameter estimated from the data. The joint probabilities of each scenario, together with the observed rank chain (SIRIUS = 1, MS-FINDER = *k*), are:

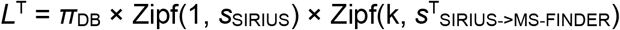

where *s*^T^_SIRIUS->MS-FINDER_ is the Zipf parameter for a true hit in the MS-FINDER ranking, conditional on the top SIRIUS hit being correct.

Similarly, the likelihoods for F and O can be approximated as follows:

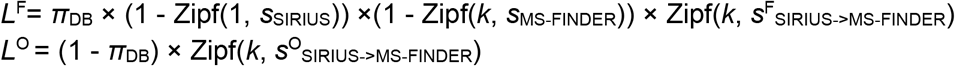

where *s*^F^_SIRIUS->MS-FINDER_ and *s*^O^_SIRIUS->MS-FINDER_ represent the Zipf parameters of the MS-FINDER rank distribution, conditional on the top SIRIUS hits being false or out of the database, respectively.

Their ratio constitutes the posterior odds (*PO*) of a correct annotation:

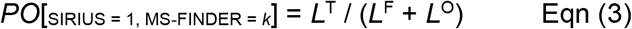

The prior odds of a correct annotation for the top SIRIUS hit, *Pr*[_SIRIUS = 1_], were

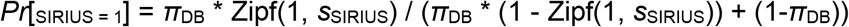

Thus, the Bayes factor of the rank chain *BF*[_SIRIUS = 1, MS-FINDER = *k*_] was defined as follows:

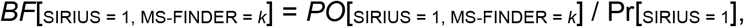

The *PO* for a chain of three ranks—such as SIRIUS = 1, MS-FINDER = *k*, and MetFrag = *l* —was calculated as follows:

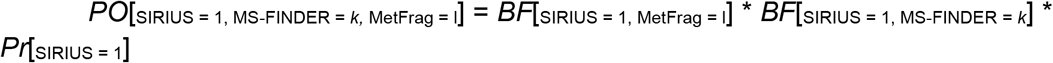

### Parameter estimation

The posterior distributions of the parameters were derived using Bayesian inference for six parameter sets: molecular formula, chemical class, and planar structure, each for the FT-MS and TOF-MS data. *π*_DB_ was set to 1.0 for molecular formula curation. The planar structure estimation from the FT-MS data is described as an example herein (Extended Data Fig. 8). All spectra annotated in Step 3 were used for the inference. First, a pre-screening step removed spectra that were intrinsically difficult to annotate (Supplementary Table 7); spectra with a SIRIUS confidence score < 0.5 were excluded. This pre-screening was essential because a small *π*_DB_ prevents *PO* from exceeding 1.0 even when a candidate is ranked first by all tools; the threshold was adjusted to keep the mode of *π*_DB_ at 1.0. For the 18,635 spectra that passed prescreening and were annotated by both MS-FINDER and SIRIUS, the distribution of the MS-FINDER ranks for the top SIRIUS hits was recorded as *D*^all^. Next, spectra with GNPS spectral library matches (spectral entropy > 0.7) were selected and their hits served as pseudo-ground truths (Supplementary Note 2). In total, 6,470 spectra whose top SIRIUS hit matched the pseudo-ground truth defined the true-hit rank distribution *D*^T^, and the 1054 unmatched spectra defined the false-hit distribution *D*^F^. To emulate the out-of-database scenario, the pseudo-ground-truth entries were removed from the SIRIUS and MS-FINDER rankings, re-ranking was performed, and the resulting MS-FINDER rank distribution among the 6,775 spectra was recorded as *D*^O^ (Extended Data Fig. 8). The observed distributions are related to the model as follows:

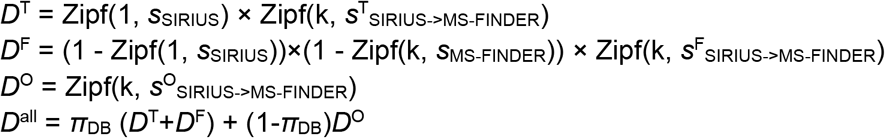

Uniform priors were assumed for *π*_DB_, *s*_SIRIUS_, *s*_MS-FINDER_, *s*^T^_SIRIUS->MS-FINDER_, *s*^F^_SIRIUS->MS-FINDER_, and *s*^O^ _SIRIUS->MS-FINDER_. Posterior distributions conditioned on the observed *D*^T^, *D*^F^, *D*^O^, and *D*^all^ were sampled using Markov chain Monte Carlo with PyMC (5,000 burn-in steps and 10,000 sampling steps)^58^ (Extended Data Fig. 9).

### Statistical decision

Because the Zipf parameters and *D*^all^ carry posterior uncertainty, the *PO* itself has a posterior distribution. The posterior median was used as the representative value of the PO, and the probability of false annotation (*FP*) was determined as follows:

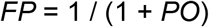

FP corresponds to the local FDR derived from the posterior error probability used in proteomics^59^.

The q-value was determined as the FDR within each dataset as follows:

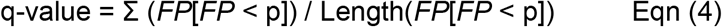

where *FP*[*FP* < p] denotes the list of *FP* values that are less than the probability *p*.

### Step 4: Curation

First, aggregated rankings were generated. The likelihood *L* for every candidate formula, class, and structure was computed using Eqn (2) based on the posterior medians of the *s* parameters. A rank of 100 was assigned to entries that were absent from the tool ranking. Candidates were sorted by normalized *L* values in descending order.

The top aggregated candidates were then evaluated. The spectra were prescreened using the criteria listed in Supplementary Table 7 and Fig. 2d. For spectra passing the prescreening, the q-value was obtained as shown in Eqn (4). If the top candidate was not ranked first by SIRIUS, MS-FINDER, or MetFrag, the PO was not evaluated. The aggregated rankings and q-values were calculated for the SIRIUS/MS-FINDER/MetFrag:PubChemLite ensemble. The SIRIUS/MS-FINDER/MetFrag:MS-FINDER DB ensemble was used in the absence of MetFrag:PubChemLite rankings.

The curation was based on the q-values under the confidence criteria described in Supplementary Note 4. We adopted the criteria of Schymanski et al.^26^ with modifications. Level 1 was not considered in the present study. Level 2a (probable structure) was assigned when both q-value < 0.05 and spectral entropy > 0.7 were achieved for an identical planar structure. Level 3 was subdivided into Levels 3-SP (tentative candidate, structure proposed; q-value < 0.05) and 3-CP (tentative candidate, class proposed; q-value < 0.05). Level 4 (unequivocal precursor ion formula; q-value < 0.01) was used for formula annotation. All the remaining spectra were assigned to Level 5 (exact mass of interest). Curation results were recorded in the COMMENT: PROPOSED_MF, COMMENT: PROPOSED_COMPOUND and COMMENT: PROPOSED_CLASS fields of each MassBank record; planar structural information for Level 2 and 3-SP records was also included in the compound fields. Supplementary Data 1 provides examples of record files.

### Metabolome analysis of a standard plasma sample

SRM 1950, Metabolites in Frozen Human Plasma developed by the National Institute of Standards and Technology, was used for metabolomic analysis^60^. Metabolites were extracted from SRM 1950 using a modified Blight–Dyer method. Briefly, 50 µL of SRM 1950 plasma was mixed with 950 µL of ice-cold methanol (−30 °C) containing the following internal standards: 10-camphorsulfonic acid and piperazine-1,4-bis(2-ethanesulfonic acid) at 1.0 nmol each, along with L-methionine sulfone, lidocaine, and 2-bromohypoxanthine at 0.2 nmol each. The mixture was then vortexed for 1 min and sonicated for 5 min. Next, the samples were incubated on ice for 5 min and centrifuged at 16,000 × *g* for 5 min at 4 °C to precipitate the proteins. The resulting supernatant (400 µL) was transferred to a 2 mL Eppendorf tube, mixed with 320 µL of Milli-Q water and 400 µL of chloroform, and vortexed for 1 min. After centrifugation at 16,000 × *g* for 5 min at 4 °C, 500 µL of the upper aqueous layer was transferred to a 1.5 mL Eppendorf tube and evaporated to dryness under vacuum using a centrifugal concentrator. The dried extract was reconstituted in 50 µL of Milli-Q water and stored at −80 °C until metabolome analysis.

LC/HRMS/MS analysis was performed using a Nexera LC system (Shimadzu Co., Kyoto, Japan) coupled with a high-performance benchtop quadrupole Orbitrap mass spectrometer (Q Exactive; Thermo Fisher Scientific, Waltham, MA, USA) equipped with an electrospray ionization source. The LC parameters were as follows: column, GL-HilicAex (2.1 × 150 mm, 5 µm particle size; Resonac Co.); injection volume, 5 µL; column temperature, 40 °C; mobile phase A, 40 mM ammonium bicarbonate in H₂O (pH 9.8); mobile phase B, acetonitrile; flow rate, 0.4 mL/min. The gradient conditions were as follows: 95% B, 0–0.5 min; 95–40% B, 0.5–15.5 min; 40–0% B, 15.5–16.5 min; 0% B, 16.5–26.5 min; 0–95% B, 26.5–27.5 min; and 95% B, 27.5–35 min.

The MS data were acquired separately for the positive and negative ion modes. The ionization conditions common to both modes were as follows: sheath gas flow rate, 50 arbitrary units; auxiliary gas flow rate, 10 arbitrary units; capillary temperature, 250 °C; S-lens level, 60; heater temperature, 400 °C. The spray voltage was set to 4,000 V and −3,000 V in positive and negative ion modes, respectively. The full-scan MS conditions were as follows: resolving power, 70,000; automatic gain control (AGC) target, 3 × 10⁶; maximum ion injection time, 200 ms; scan range, m/z 70–1,050. The data-dependent MS/MS (dd-MS/MS) conditions were as follows: resolving power, 17,500; AGC target, 1 × 10⁶; maximum ion injection time, 80 ms; loop count, 20; isolation width, ±0.2 Da; fixed first mass, m/z 50; normalized collision energy, 20. The intensity threshold for precursor ions and the dynamic exclusion time for dd-MS/MS were set to 1 × 10³ and 6 s, respectively.

### Metabolome analysis of cancer cell lines

HCT116 colorectal cancer cells were obtained from the American Type Culture Collection (ATCC; Manassas, VA, USA), and HCT116 IDH1 (R132H/+) cells were obtained from Horizon Discovery Ltd. (Cambridge, UK). The culture experiments were conducted as described previously^61^. Briefly, 6.0 × 105 cells were seeded in 5 mL of Roswell Park Memorial Institute 1640 (RPMI 1640) medium (Wako) supplemented with 10% fetal bovine serum and antibiotics (100 U/mL penicillin, 100 μg/mL streptomycin, and 0.25 μg/mL amphotericin B; Nacalai Tesque) in 60-mm-diameter plates and pre-cultured for 24 h at 37 °C under 5% CO_2_. The cells were then washed twice with D-PBS (Nacalai Tesque). Next, the culture medium was replaced with Advanced RPMI 1640 medium (Thermo Fisher Scientific) containing 2 mmol/L glutamine (Nacalai Tesque) and antibiotics, and the cells were cultured for 2 days.

The intracellular metabolites were extracted using the methanol/water/chloroform method^62^. After rapid removal of the medium and rinsing with 5 mL of 5% (w/v) mannitol (Wako) in Milli-Q water (Merck Millipore), cellular metabolism was rapidly quenched by adding 700 µL of pre-cooled methanol containing 1 μmol/L 10-camphorsulfonic acid (Wako), L-tryptophan-13C11-15N2 (Nippon Sanso), and L-methionine sulfone (Alfa Aesar). The cells and solutions were collected by scraping. The cell lysate was then transferred to a new tube and sonicated for 5 min. The supernatant (400 μL) was mixed vigorously with 160 μL of methanol, 280 μL of chloroform, and 160 μL of Milli-Q water using a vortex mixer for 1 min, followed by 5 min of sonication. After centrifugation (4 °C, 16,000 × *g*, 5 min), 800 μL of the supernatant was transferred to a new tube, and 220 μL of chloroform and 220 μL of Milli-Q water were added. After vortexing and centrifugation (4 °C, 16,000 × *g*, 3 min), 500 µL of the upper aqueous layer was collected and dried in a vacuum centrifugal concentrator (CentriVap, LABCONCO) at room temperature.

The dried samples were resuspended in 125 μL of 50% (v/v) acetonitrile (Wako) in Milli-Q water and subjected to metabolomic analysis using an Agilent 1290 Infinity LC system (Agilent Technologies) coupled with a Q Exactive Orbitrap MS system (Thermo Fisher Scientific), following a previously described method with some modifications^63^. Briefly, 1 µL of the sample was injected into an InfinityLab Poroshell 120 HILIC-Z column (2.1 mm × 150 mm, 2.7 µm; Agilent Technologies). The flow rate was 0.25 mL/min, and the column temperature was set to 40 °C. The mobile phase consisted of 20 mmol/L ammonium formate + 0.25% (v/v) formic acid (A) and 20 mmol/L ammonium formate + 0.25% formic acid in 90% (v/v) acetonitrile (B). The separation was performed using the following gradient: 0 min, 100% B; 15 min, 70% B; 20 min, 10% B; and 23 min, 10% B. The column was equilibrated with the initial mobile phase composition for 7 min before the next injection. The analysis was performed using DDA in positive ion mode with the following source parameters: auxiliary gas temperature, 300 °C; auxiliary gas flow rate, 10 (arbitrary units); spray voltage, 3.5 kV; capillary temperature, 250 °C; sheath gas flow rate, 40 (arbitrary units); and S-lens, 35 (arbitrary units).

The acquisition parameters were as follows: MS1 and MS2 mass range, m/z 50–750; MS1 event time, 200 ms; MS1 resolution, 70,000; MS1 AGC target, 3.0 × 106; MS1 maximum ion injection time, 200 ms; MS2 resolution, 17,500; MS2 AGC target, 1.0 × 105; MS2 maximum ion injection time, 50 ms; number of MS2 events, 5; MS2 target range, m/z 200–2,000; dynamic exclusion duration, 10 s; NCE, 30; CID gas, N2. Data were processed using MS-DIAL version 5^64^ with default settings for all parameters. Metabolite annotation was performed using MS-DIAL and a spectral library (MSMS-Public_experimentspectra-pos-VS19). msp) provided on the official MS-DIAL website (https://systemsomicslab.github.io/compms/msdial/main.html).

## Data availability

All the outputs of the ShinMassBank reprocessing pipeline, including the MassBank record, MSP, and mzTab-M files, are available at https://github.com/Shin-MassBank/MassBank-Human. The raw data files of the two DDA metabolome datasets acquired in this study have been deposited in MB-POST (MPST000004 and MPST000223). Spectrum files in .ms format for SIRIUS searches (Supplementary Data 2) have been deposited in Zenodo (https://doi.org/10.5281/zenodo.22968507).

## Code availability

The Python scripts for the ShinMassBank reprocessing pipeline are available at https://github.com/fumiomatsuda/ShinMassBank-reprocessing-pipeline.

## Acknowledgements

The authors thank all the data submitters, data repository developers, and tool developers for their contributions. The authors are also grateful for the support provided by MassBank developers.

## Contributions

F.M. A.H., Y.I., S.O., H.T., S.N. and E.L.S. conceptualized the study. F.M., R.H. Y.T., A.C.Y. and M.T. performed the data analysis. F.M., R.H. and M.T. drafted the paper. F.M., R.H., and T.T. processed the data. F.M. and A.H. supervised the study. R.H. and M.T. collected metabolome data. F.M. A.H., Y.I., S.O., H.T., S.N. and E.L.S. discussed the results, critically reviewed the text and approved the final paper.

---

**Extended Data Fig. 1.**
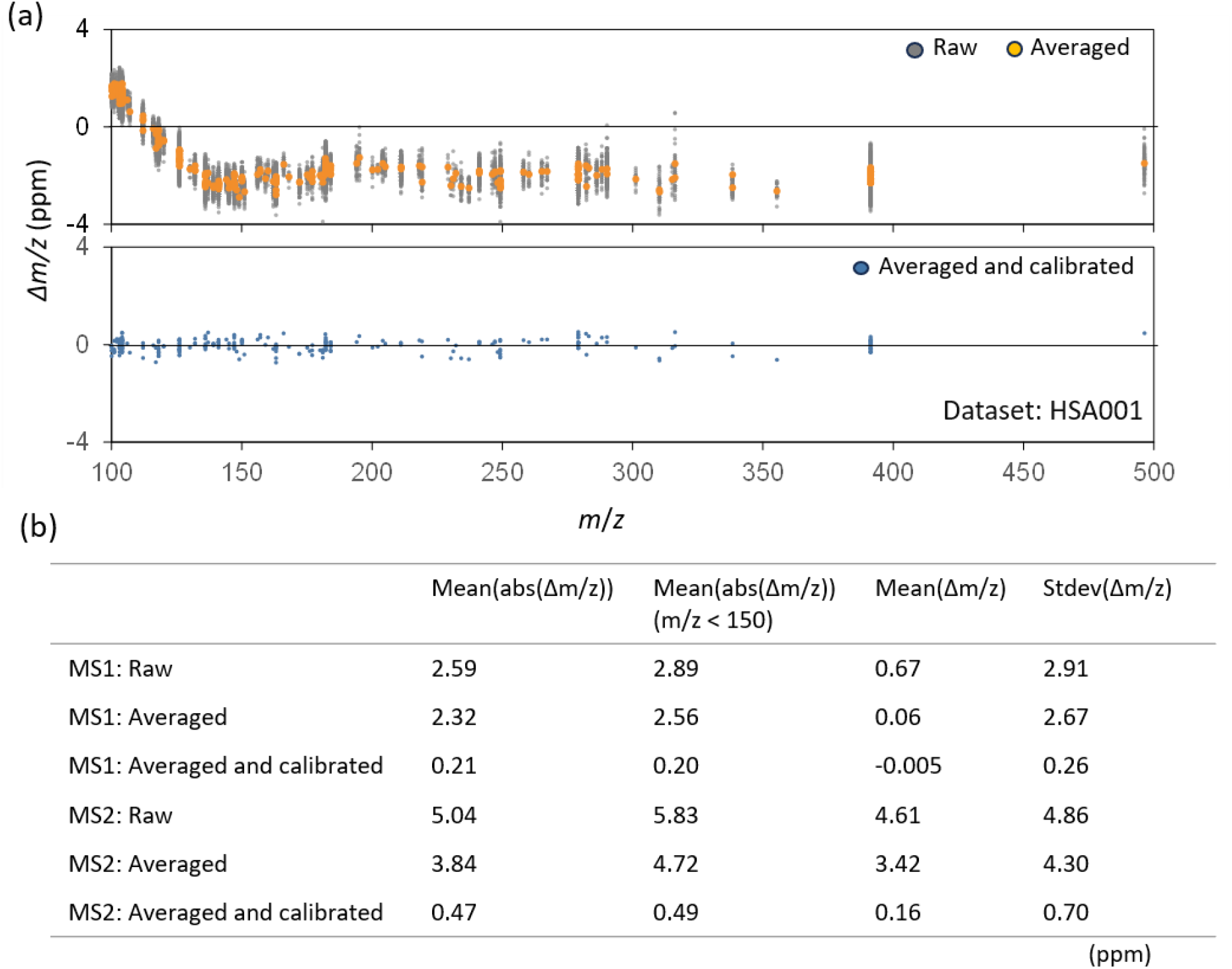
Improvement of *m*/*z* accuracy by spectral averaging and data-dependent calibration. (a) The x-axis shows measured *m*/*z* values in MS1 spectra of the HSA001 dataset; the y-axis shows the deviation of measured from theoretical *m*/*z* values (Δ*m*/*z*). (b) Summary of improvements in data quality following averaging and calibration.

**Extended Data Fig. 2.**
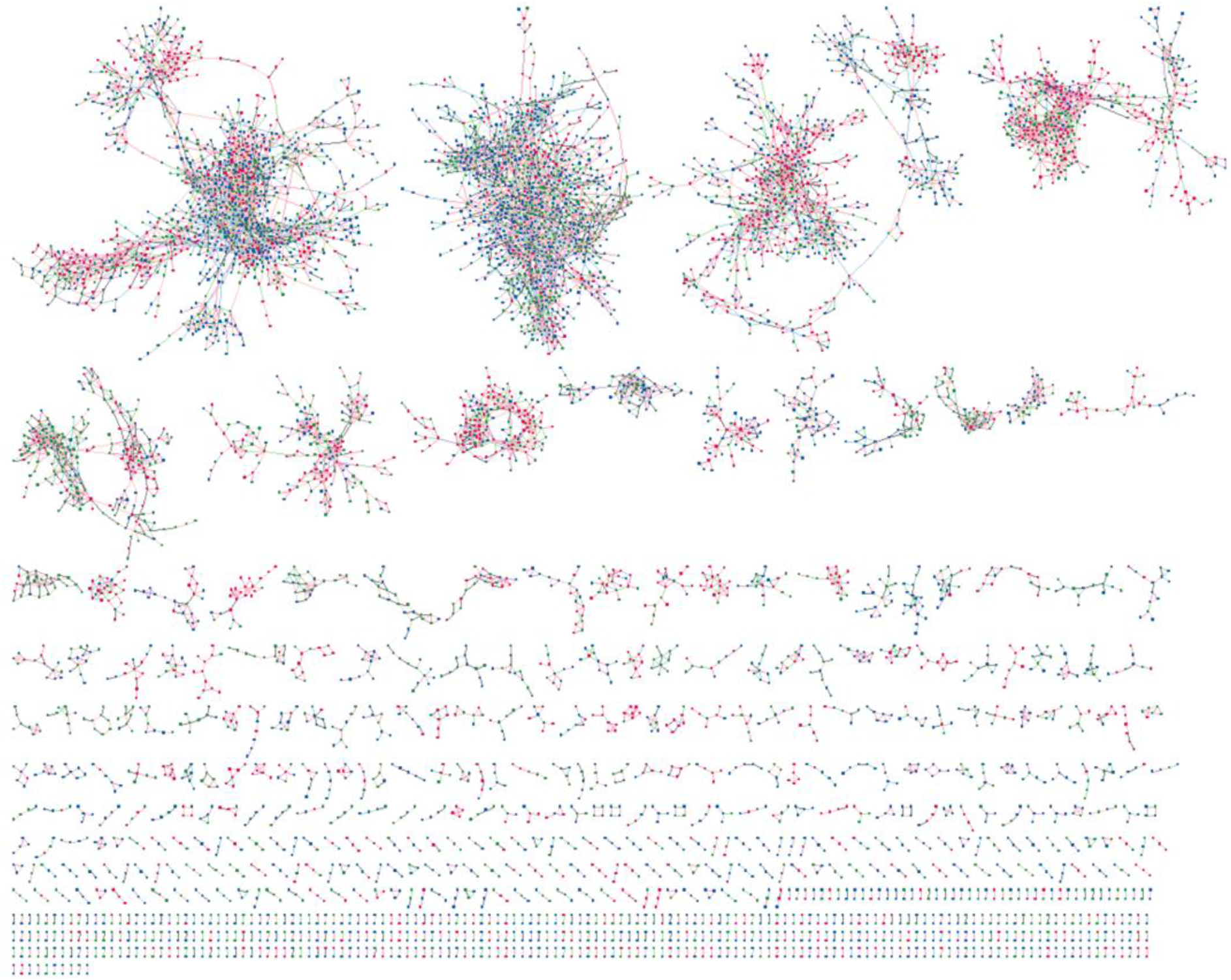
Molecular Network of MassBank Human Spectra. Spectra curated at Levels 2a or 3-SP (planar structure, red), 3-CP (class, green), and 4 (molecular formula, blue) were used for network construction.

**Extended Data Fig. 3.**
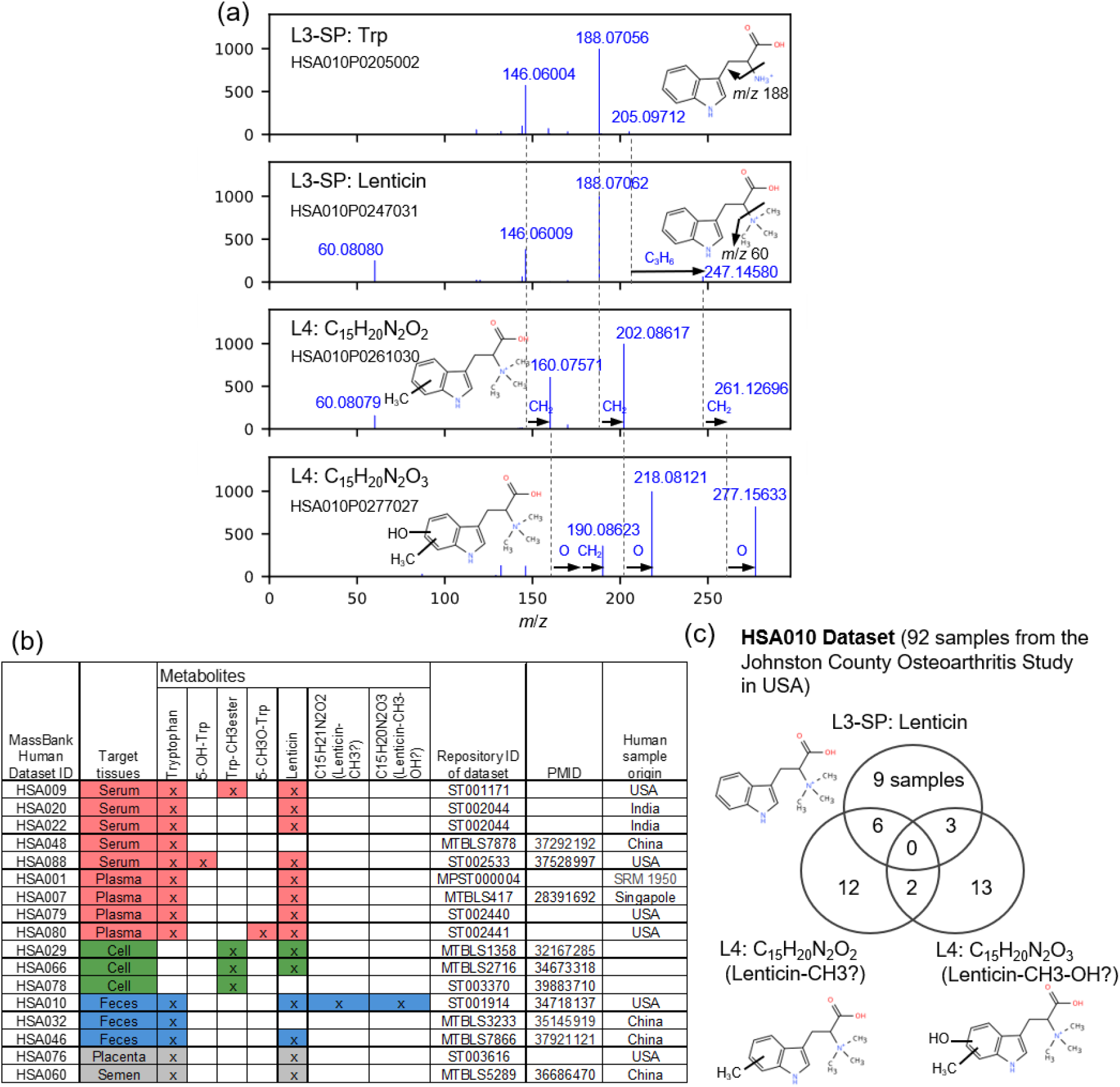
Biological distribution and spectral characteristics of lenticin and its derivatives. Lenticin (**1**) is a trimethylated tryptophan-derived betaine originating from lentils. (a) MS2 spectral comparison of tryptophan (Trp), lenticin, and their derivatives (C_15_H_20_N_2_O_2_ and C_15_H_20_N_2_O_3_). The arrows indicate mass shifts corresponding to methylation (+CH_2_) and hydroxylation (+O), supporting their structural relationships. (b) Detection patterns across datasets. Matrix shows the presence (red, serum; green, cells; blue, feces) of Trp, lenticin, and related analogs across 76 datasets. The presence/absence pattern highlights differences in dietary and biological contexts. (c) Interindividual variations in the HSA010 dataset. The Venn diagram shows the detection of lenticin and its methylated/hydroxylated derivatives in 92 samples from a US fecal cohort. The varying overlap suggests individual differences in the gut microbiota-mediated transformations.

**Extended Data Fig. 4.**
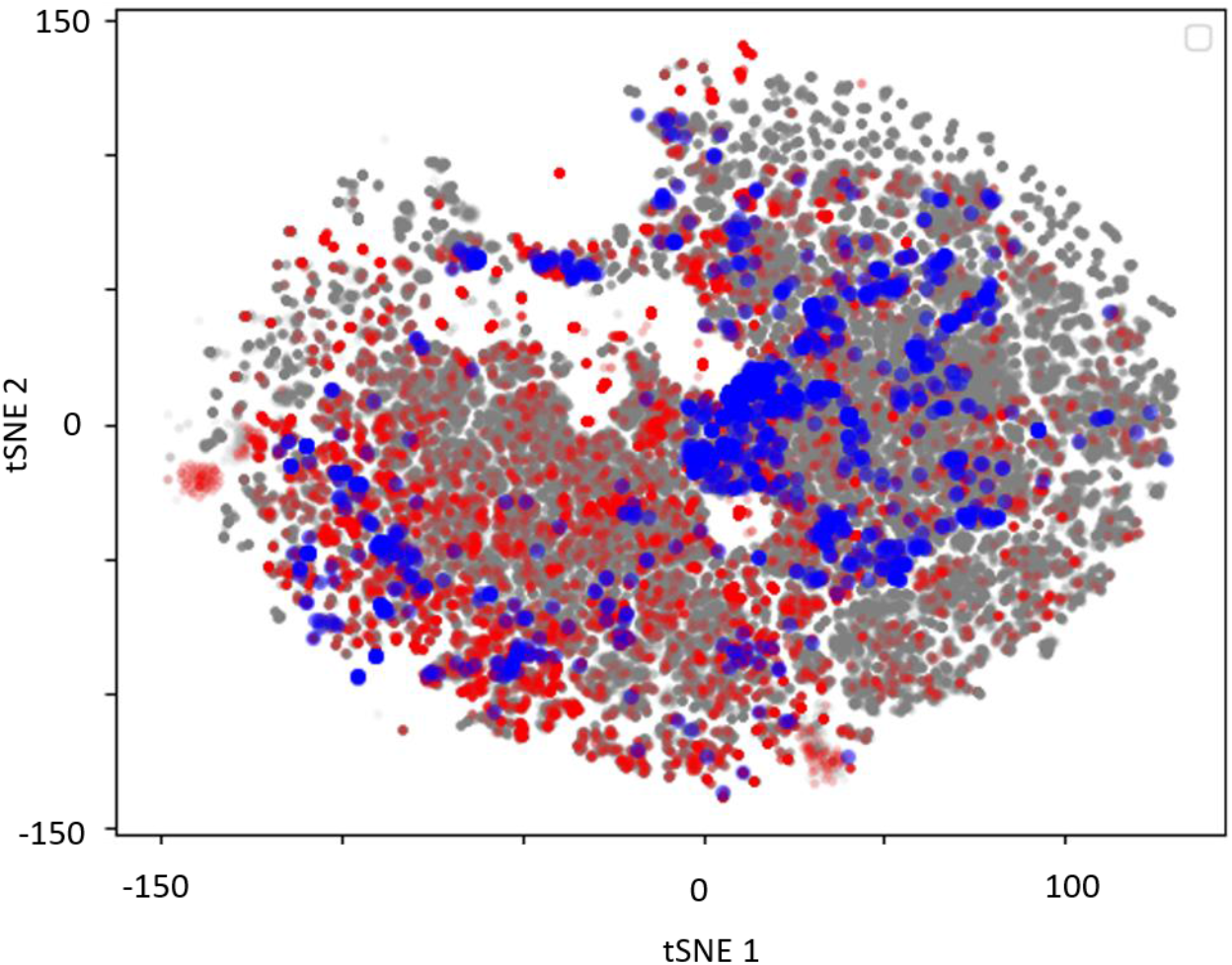
Metabolite space of MassBank Human. Metabolites in the MS-FINDER compound database (554,833 compounds) are projected using t-SNE with their fingerprints (MACCS keys). Metabolites in the Human Metabolome Database (HMDB) and those curated at Levels 2a or 3-SP in MassBank Human are highlighted in red and blue, respectively.

**Extended Data Fig. 5.**
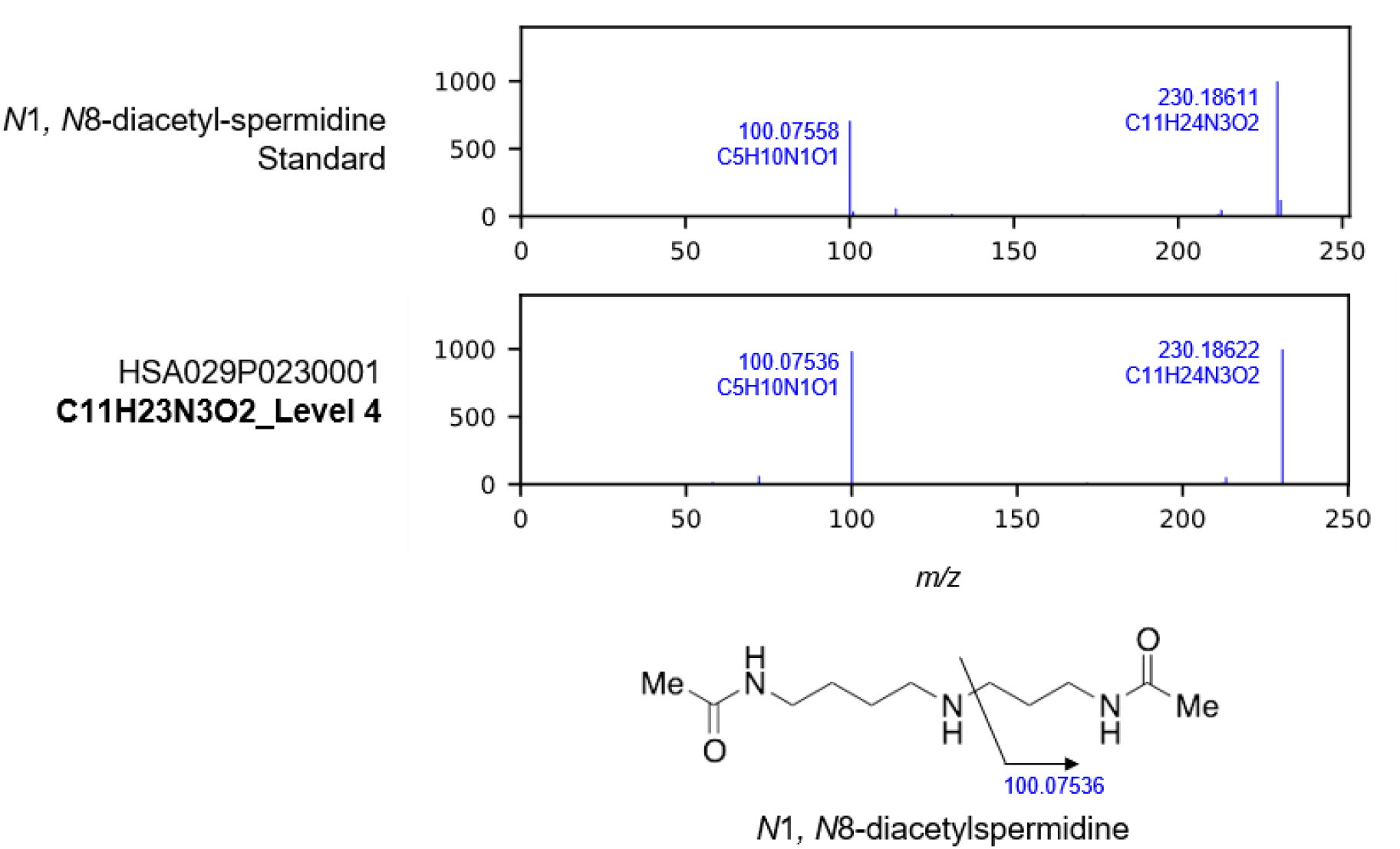
Validation of the *N*1,*N*8-diacetylspermidine annotation using an authentic standard. To confirm the identity of *N*1,*N*8-diacetylspermidine in *IDH1*-mutant colorectal cancer cells, the MS2 spectrum was compared to that of a high purity chemical standard. The top and bottom panels show the standard and MassBank Human spectrum, respectively, demonstrating matching fragment ions and relative intensities.

**Extended Data Fig. 6.**
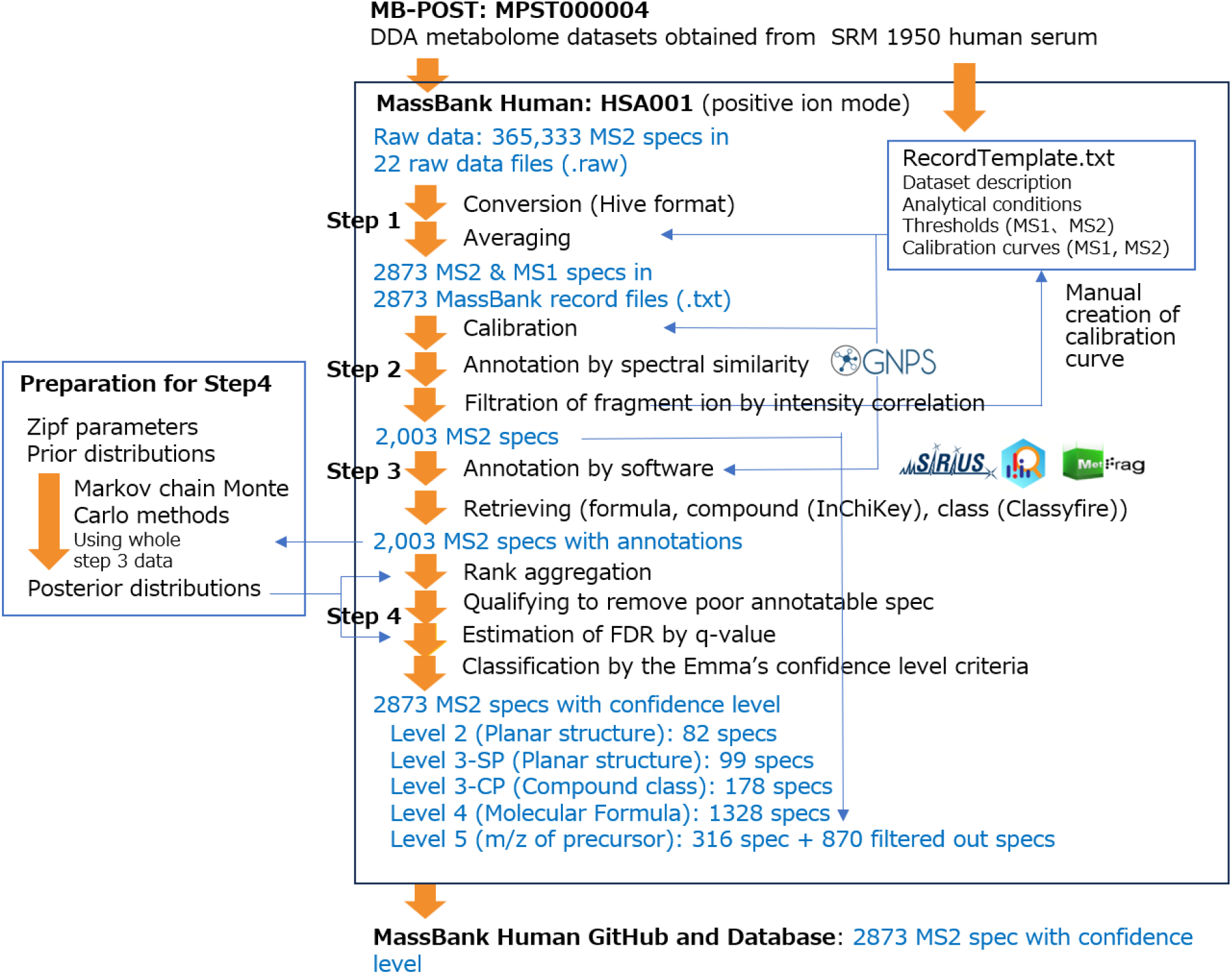
Overview of the reprocessing pipeline and an example using the HSA001 dataset.

**Extended Data Fig. 7.**
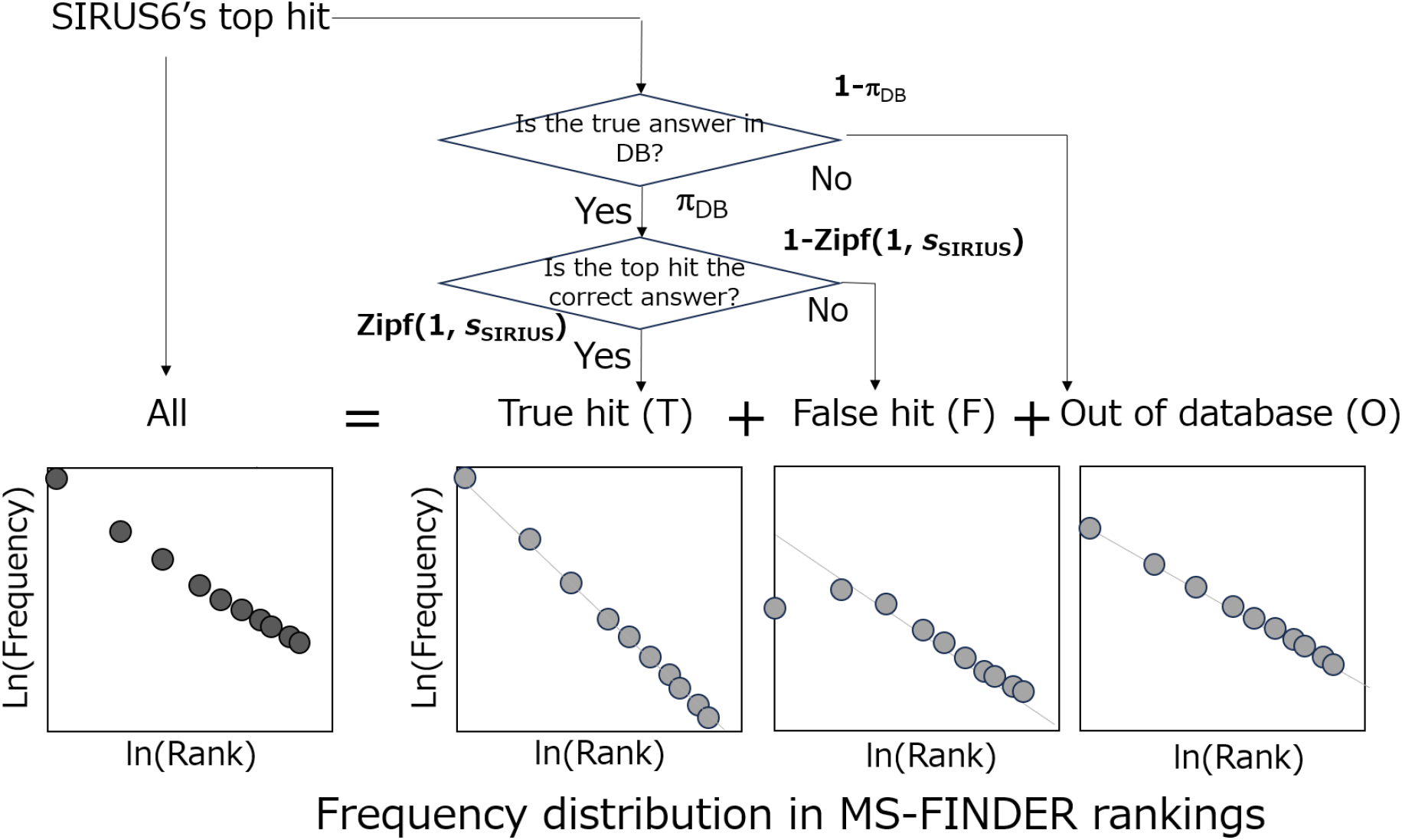
Conceptual framework of the rank-chain model in ZBRA. The ZBRA framework integrates rankings from multiple tools by modeling how the top hit from one software tool behaves in the ranking of another. The total observed frequency distribution ("All") is modeled as a mixture of three scenarios: true hit (T; the top hit is correct), false hit (F; a wrong candidate is ranked first), and out of database (O; the true compound is not in the search space). Each scenario followed the Zipf distribution with specific parameters.

**Extended Data Fig. 8.**
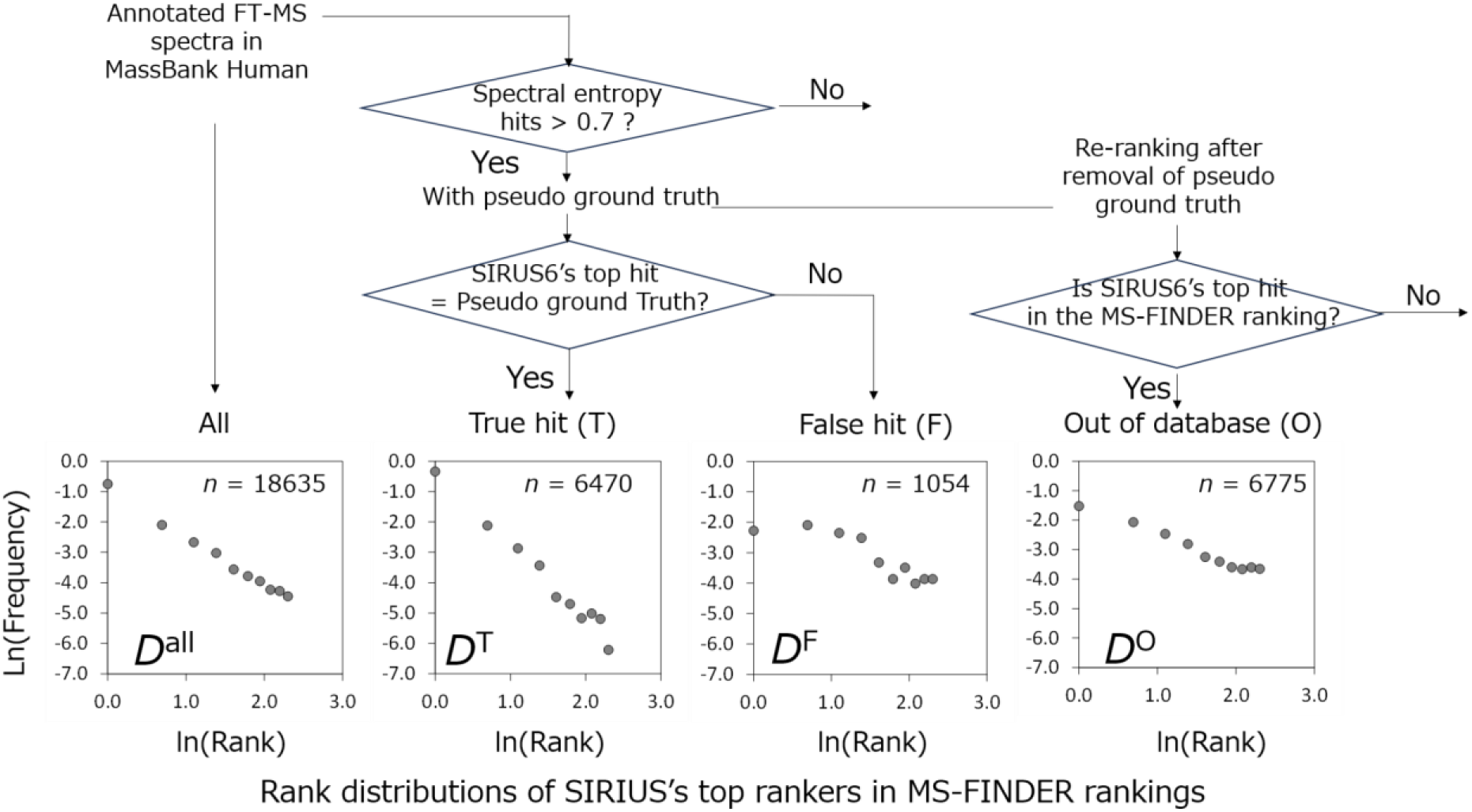
Empirical rank distributions used for Bayesian parameter estimation. To calibrate the ZBRA model, the pipeline used spectral library matches as pseudo-ground truths to estimate the Zipf parameters. The log–log plots show the rank distribution of the top SIRIUS hits within the MS-FINDER rankings for the FT-MS planar structures. The flowchart shows the classification procedure used in this study. Distributions are shown for all spectra (*D*_all_), confirmed true hits (*D*^T^), confirmed false hits (*D*^F^), and a simulated out-of-database scenario (*D*^O^) in which ground truths were intentionally removed.

**Extended Data Fig. 9.**
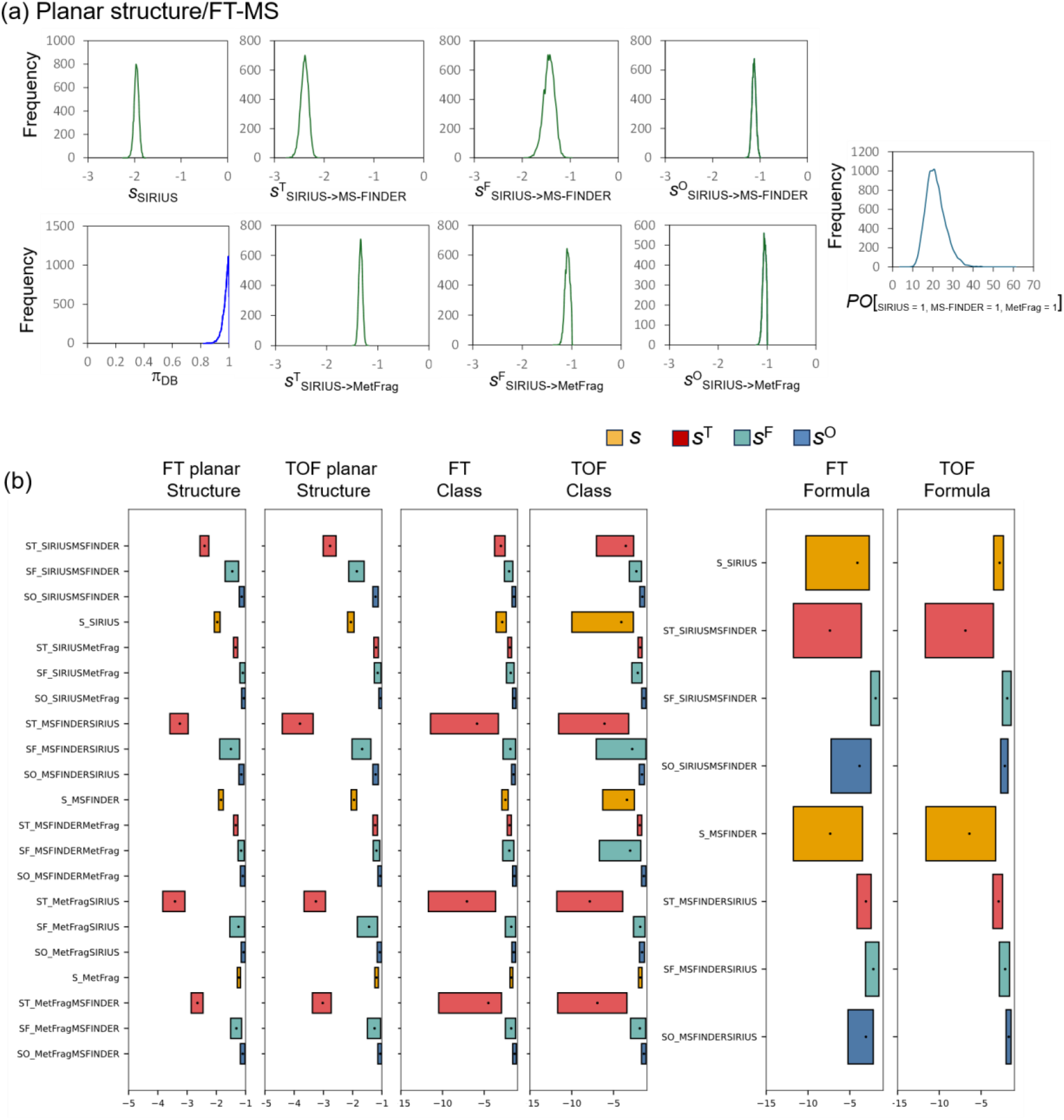
Posterior distributions of parameters and posterior odds (*PO*). The ZBRA framework estimates the posterior distributions of the parameters and *PO* using Markov chain Monte Carlo (MCMC) methods. (a) Posterior distributions of the Zipf parameters and *PO*. (b) Summary of posterior distributions of the estimated parameters. Box plots summarize the ranges of Zipf parameters (*s*, *s*^T^, *s*^F^, and *s*^O^) derived for different annotation tiers (planar structure, chemical class, and molecular formula) and instrument types (FT vs. TOF).

## Supplementary information

Supplementary Information (pdf)

Supplementary Notes 1-6

Supplementary Figures 1-5

Supplementary Table (Excel)

Supplementary Tables 1–11.

Supplementary Data (zipped CSV)

Supplementary Data 1, 3-9.

Supplementary Data 2 is available from Zenodo (DOI: 10.5281/zenodo.22968507).

## References

1 Johnson, C. H., Ivanisevic, J. & Siuzdak, G. Metabolomics: beyond biomarkers and towards mechanisms. Nat. Rev. Mol. Cell Biol. 17, 451–459 (2016).

2 Lai, Y. et al. High-Resolution Mass Spectrometry for Human Exposomics: Expanding Chemical Space Coverage. Environ. Sci. Technol. 58, 12784–12822 (2024).

3 Krautkramer, K. A., Fan, J. & Backhed, F. Gut microbial metabolites as multi-kingdom intermediates. Nat. Rev. Microbiol. 19, 77–94 (2021).

4 da Silva, R. R., Dorrestein, P. C. & Quinn, R. A. Illuminating the dark matter in metabolomics. Proc. Natl. Acad. Sci. U. S. A. 112, 12549–12550 (2015).

5 Kind, T. et al. Identification of small molecules using accurate mass MS/MS search. Mass Spectrom. Rev. 37, 513–532 (2018).

6 Scheubert, K. et al. Significance estimation for large scale metabolomics annotations by spectral matching. Nat. Commun. 8, 1494 (2017).

7 Wang, M. et al. Mass spectrometry searches using MASST. Nat. Biotechnol. 38, 23–26 (2020).

8 Matsuda, F. et al. MS/MS spectral tag (MS2T)-based annotation of non-targeted profile of plant secondary metabolites. Plant J. 57, 555–577 (2009).

9 Kind, T. et al. FiehnLib: mass spectral and retention index libraries for metabolomics based on quadrupole and time-of-flight gas chromatography/mass spectrometry. Anal. Chem. 81, 10038–10048 (2009).

10 Fiehn, O. et al. Metabolite profiling for plant functional genomics. Nat. Biotechnol. 18, 1157–1161 (2000).

11 Sud, M. et al. Metabolomics Workbench: An international repository for metabolomics data and metadata, metabolite standards, protocols, tutorials and training, and analysis tools. Nucleic Acids Res. 44, D463–470 (2016).

12 Yurekten, O. et al. MetaboLights: open data repository for metabolomics. Nucleic Acids Res. 52, D640–D646 (2024).

13 Nothias, L. F. et al. Feature-based molecular networking in the GNPS analysis environment. Nat. Methods 17, 905–908 (2020).

14 Takahashi, Y. et al. Reviving data potential horizontal line-The reanalysis-driven metabolomics data repository MB-POST. Anal. Chem. (2026).

15 Hoffmann, N. et al. mzTab-M: A data standard for sharing quantitative results in mass spectrometry metabolomics. Anal. Chem. 91, 3302–3310 (2019).

16 Matsuda, F. Data processing of Product ion spectra: Redundancy of product ion spectra of small molecules in data-dependent acquisition dataset. Mass Spectrom (Tokyo) 12, A0138 (2023).

17 Frank, A. M. et al. Clustering millions of tandem mass spectra. J. Proteome Res. 7, 113–122 (2008).

18 Wang, M. et al. Sharing and community curation of mass spectrometry data with Global Natural Products Social Molecular Networking. Nat. Biotechnol. 34, 828–837 (2016).

19 Zhang, N. H. & Deutsch, E. W. SpectiCal: m/z calibration of MS2 peptide spectra using known low mass ions. J. Proteome Res. 23, 1519–1530 (2024).

20 Stravs, M. A., Schymanski, E. L., Singer, H. P. & Hollender, J. Automatic recalibration and processing of tandem mass spectra using formula annotation. J. Mass Spectrom. 48, 89–99 (2013).

21 Duhrkop, K. et al. SIRIUS 4: a rapid tool for turning tandem mass spectra into metabolite structure information. Nat. Methods 16, 299–302 (2019).

22 Tsugawa, H. et al. Hydrogen rearrangement rules: Computational MS/MS fragmentation and structure elucidation using MS-FINDER software. Anal. Chem. 88, 7946–7958 (2016).

23 Ruttkies, C., Neumann, S. & Posch, S. Improving MetFrag with statistical learning of fragment annotations. BMC Bioinformatics 20, 376 (2019).

24 Koistinen, V. et al. Towards a Rosetta stone for metabolomics: recommendations to overcome inconsistent metabolite nomenclature. Nat Metab 5, 351–354 (2023).

25 Newman, M. Power laws, Pareto distributions and Zipf’s law. Contemporary Physics 46, 323–351 (2005).

26 Schymanski, E. L. et al. Identifying small molecules via high resolution mass spectrometry: communicating confidence. Environ. Sci. Technol. 48, 2097–2098 (2014).

27 Djoumbou Feunang, Y., et al. ClassyFire: automated chemical classification with a comprehensive, computable taxonomy. J Cheminform 8, 61 (2016).

28 Vos, M. E., Meima, M. Y. & Houben, G. F. Cleaning and using FooDB: a method for extracting compound information and applying FooDB in dietary cohort studies. MethodsX 15, 103691 (2025).

29 Kruger, R. et al. MiMeDB 2.0: the human microbial metabolome database for 2026. Nucleic Acids Res. 54, D623–D634 (2026).

30 Elapavalore, A. et al. PubChemLite Plus Collision coss section (CCS) values for enhanced interpretation of nontarget environmental data. Environ Sci Technol Lett 12, 166–174 (2025).

31 Vermeulen, R., Schymanski, E. L., Barabasi, A. L. & Miller, G. W. The exposome and health: Where chemistry meets biology. Science 367, 392–396 (2020).

32 Wikoff, W. R. et al. Metabolomics analysis reveals large effects of gut microflora on mammalian blood metabolites. Proc. Natl. Acad. Sci. U. S. A. 106, 3698–3703 (2009).

33 van Kessel, S. P. et al. Gut bacterial tyrosine decarboxylases restrict levels of levodopa in the treatment of Parkinson’s disease. Nat. Commun. 10, 310 (2019).

34 Tomas-Barberan, F. A. et al. Urolithins, the rescue of "old" metabolites to understand a "new" concept: Metabotypes as a nexus among phenolic metabolism, microbiota dysbiosis, and host health status. Mol. Nutr. Food Res. 61 (2017).

35 Peisl, B. Y. L., Schymanski, E. L. & Wilmes, P. Dark matter in host-microbiome metabolomics: Tackling the unknowns-A review. Anal. Chim. Acta 1037, 13–27 (2018).

36 Yu, J. S. et al. Tandem Mass Spectrometry Molecular Networking as a Powerful and Efficient Tool for Drug Metabolism Studies. Anal. Chem. 94, 1456–1464 (2022).

37 Garcia-Aloy, M. et al. Discovery of intake biomarkers of lentils, chickpeas, and white beans by untargeted LC-MS metabolomics in serum and urine. Mol. Nutr. Food Res. 64, e1901137 (2020).

38 Rushing, B. R. et al. Fecal metabolomics reveals products of dysregulated proteolysis and altered microbial metabolism in obesity-related osteoarthritis. Osteoarthritis Cartilage 30, 81–91 (2022).

39 Wishart, D. S. et al. HMDB 5.0: the Human metabolome database for 2022. Nucleic Acids Res. 50, D622–D631 (2022).

40 Dang, L. et al. Cancer-associated IDH1 mutations produce 2-hydroxyglutarate. Nature 462, 739–744 (2009).

41 Umemori, Y. et al. Evaluating the utility of N1,N12-diacetylspermine and N1,N8-diacetylspermidine in urine as tumor markers for breast and colorectal cancers. Clin. Chim. Acta 411, 1894-1899 (2010).

42 Neumann, S. et al. MassBank: an open and FAIR mass spectral data resource. Nucleic Acids Res. 54, D601–D606 (2026).

43 Casero, R. A., Jr., Murray Stewart, T. & Pegg, A. E. Polyamine metabolism and cancer: treatments, challenges and opportunities. Nat. Rev. Cancer 18, 681–695 (2018).

44 Gerlich, M. & Neumann, S. MetFusion: integration of compound identification strategies. J. Mass Spectrom. 48, 291–298 (2013).

45 Schymanski, E. L. et al. Critical assessment of small molecule identification 2016: automated methods. J Cheminform 9, 22 (2017).

46 Kolde, R., Laur, S., Adler, P. & Vilo, J. Robust rank aggregation for gene list integration and meta-analysis. Bioinformatics 28, 573–580 (2012).

47 Palmer, A. et al. FDR-controlled metabolite annotation for high-resolution imaging mass spectrometry. Nat. Methods 14, 57–60 (2017).

48 Hoffmann, M. A. et al. High-confidence structural annotation of metabolites absent from spectral libraries. Nat. Biotechnol. 40, 411–421 (2022).

49 Shteynberg, D. et al. iProphet: multi-level integrative analysis of shotgun proteomic data improves peptide and protein identification rates and error estimates. Mol. Cell. Proteomics 10, M111 007690 (2011).

50 Giera, M., Aisporna, A., Uritboonthai, W. & Siuzdak, G. The hidden impact of in-source fragmentation in metabolic and chemical mass spectrometry data interpretation. Nat Metab 6, 1647–1648 (2024).

51 Ludwig, M., Fleischauer, M., Duhrkop, K., Hoffmann, M. A. & Bocker, S. De novo molecular formula annotation and structure elucidation uing SIRIUS 4. Methods Mol. Biol. 2104, 185–207 (2020).

52 Schymanski, E. L. et al. Empowering large chemical knowledge bases for exposomics: PubChemLite meets MetFrag. J Cheminform 13, 19 (2021).

53 Horai, H. et al. MassBank: a public repository for sharing mass spectral data for life sciences. J. Mass Spectrom. 45, 703–714 (2010).

54 Li, Y. et al. Spectral entropy outperforms MS/MS dot product similarity for small-molecule compound identification. Nat. Methods 18, 1524–1531 (2021).

55 Watrous, J. et al. Mass spectral molecular networking of living microbial colonies. Proc. Natl. Acad. Sci. U. S. A. 109, E1743–1752 (2012).

56 Kohl, M., Wiese, S. & Warscheid, B. Cytoscape: software for visualization and analysis of biological networks. Methods Mol. Biol. 696, 291–303 (2011).

57 Matsuda, F., Komori, S., Yamada, Y., Hara, D. & Okahashi, N. Data processing of product ion spectra: Quality improvement by averaging multiple similar spectra of small molecules. Mass Spectrom (Tokyo) 11, A0106 (2022).

58 Salvatier, J., Wiecki, T. V. & Fonnesbeck, C. Probabilistic programming in Python using PyMC3. PeerJ Computer Science 2, e55 (2016).

59 Kall, L., Storey, J. D., MacCoss, M. J. & Noble, W. S. Assigning significance to peptides identified by tandem mass spectrometry using decoy databases. J. Proteome Res. 7, 29–34 (2008).

60 Phinney, K. W. et al. Development of a standard reference material for metabolomics research. Anal. Chem. 85, 11732–11738 (2013).

61 Hayasaka, R. et al. Metabolomics of small extracellular vesicles derived from isocitrate dehydrogenase 1-mutant HCT116 cells collected by semi-automated size exclusion chromatography. Front Mol Biosci 9, 1049402 (2022).

62 Hayasaka, R. et al. Metabolomic analysis of small extracellular vesicles derived from pancreatic cancer cells cultured under normoxia and hypoxia. Metabolites 11 (2021).

63 Suzuki, Y. et al. Comparative metabolomics of small molecules specifically expressed in the dorsal or ventral marginal zones in vertebrate gastrula. Metabolites 12 (2022).

64 Tsugawa, H. et al. MS-DIAL: data-independent MS/MS deconvolution for comprehensive metabolome analysis. Nat. Methods 12, 523–526 (2015).

