## Supplementary information for "MassBank Human: The annotated human metabolome"

**Contents**

**Supplementary Note 1.** Detailed procedure of spectral preprocessing
**Supplementary Note 2.** FDR estimation of spectral search results
**Supplementary Note 3.** Rank aggregation using the Zipf distribution
**Supplementary Note 4.** Extension of annotation confidence level criteria
**Supplementary Note 5.** Assessment of the performance of Zipf–Bayesian rank aggregation
**Supplementary Note 6.** Structural annotation of tocopherols using the inter-dataset molecular network

**Supplementary Figure 1.** An example of preprocessing
**Supplementary Figure 2.** Frequency distribution of correct hits in the planar structure ranking by MS-FINDER
**Supplementary Figure 3**. Frequency distribution of correct hits in the molecular formula, class, and planar structure rankings by MS-FINDER, SIRIUS, MetFrag, and their aggregates
**Supplementary Figure 4.** Evaluation of the FDR through comparison with GNPS library search results
**Supplementary Figure 5.** Structural elucidation of α-tocopherol derivatives (vitamin E) using the inter-dataset molecular network

### Supplementary Notes

**Supplementary Note 1.** Detailed procedure of spectral preprocessing

In this section, we explain the spectral averaging performed in this study using an example. The HSA001 dataset consists of 22 data files acquired by DDA on a Q Exactive mass spectrometer from a standard reference material (SRM 1950 human plasma). It contains a total of 365,333 MS2 spectra.

The MS/MS spectrum in the 16_posi.raw file of HSA001 (retention time 9.020 min; MS2 scan no. 4,471 acquired from precursor ion *m/z* 205.097162; Supplementary Figure 1a and Supplementary Table 8) was identified as tryptophan based on its chromatographic and mass spectral behavior[1]. A comparison revealed that fragment ions reported in a previous tryptophan fragmentation study[2] matched 9 of the 21 fragment ions in the raw spectra (Supplementary Table 8).

Within HSA001, 33 similar spectra acquired from the same precursor ion were present (|Δretention time| < 1.5 min, precursor ion |Δ*m/z*| < ms1_mz_raw_thres = 1.5 ppm, and product ion spectrum cosine product score > 0.9). These were processed to generate an averaged and recalibrated spectrum (Supplementary Figure 1b). Only ions observed in at least 70% (≥24) of the 33 spectra were retained during averaging. The resulting spectrum retained 10 of the 21 fragment ions, as shown in Supplementary Figure 1a and Supplementary Table 8. These fragment ions represented the signals reproducibly observed more than 24 raw spectra. Of these, eight matched the tryptophan fragmentation study. One matching ion (*m/z* 130.06506) was removed during averaging, likely because it was the product of a minor fragmentation pathway[2]. A comparison between the theoretical and observed *m/z* values of the MS2 product ions confirmed that averaging and recalibration improved mass accuracy (Supplementary Figure 1d).

Next, the correlation between the product ion and precursor ion intensities across the 33 raw spectra was evaluated for the 10 product ions shown in Supplementary Figure 1b. Two non-matching ions, *m/z* 87.0440763 and 149.0233415, showed low correlation coefficients ($r$ = 0.13 and 0.37, respectively; Supplementary Figure 1e, f), suggesting that they were interference-derived signals (Supplementary Table 8). Conversely, eight matching ions displayed significant positive correlations ($r$ = 0.98 at *m/z* 188.07064 (Supplementary Figure 1g). The *m/z* values of the final retained fragment ions matched the theoretical values with sub-ppm accuracy (Supplementary Table 8). These results demonstrate that averaging, recalibration, and filtering markedly enhances the quality of MS2 spectra. This spectrum was assigned the ID "HSA001P0205016."

**Supplementary Note 2.** FDR estimation of spectral search results

In this study, spectral similarity search results were used as the pseudo-ground truth to construct an annotation methodology. First, we estimated the FDR of the spectral search results to verify whether the similarity search results could serve as a pseudo-ground truth. MassBank spectra were searched against the GNPS spectral library[3] to evaluate the FDR of the search results. FT-MS and TOF-MS spectra were extracted from MassBank to create evaluation datasets (FT-MS: 3,234 spectra; TOF-MS: 12,464 spectra). Each spectrum in these datasets was queried against the GNPS spectral library (see Methods). Top-scoring hits with a spectral entropy[4] score > 0.7 were considered hits, ignoring self-hits.

The molecular formulae of the query and hit compounds were compared (Supplementary Table 9). For the FT dataset, 2,037 spectra had hits at a score threshold of 0.7, all of which matched the query (FDR = 0%). For the TOF dataset, 7,874 spectra had hits; with 60 mismatches were detected (FDR = 0.0076). Based on these results, the molecular formulae of compounds hit in spectral searches at a threshold of 0.7 were considered valid as pseudo-ground truths with an FDR < 1%.

The planar structures of the query and hit compounds were compared (Supplementary Table 9). Identity was determined by matching the first block of the InChIKey. At a similarity score threshold of 0.7, the FDR values for the FT and TOF datasets were 0.0363 (74/2,038) and 0.0924 (728/7,880), respectively. Detailed inspection of the non-matching cases revealed that most were structural isomers with similar chemical structures (Supplementary Data 8).

Finally, ChemOnt classifications of the query and hit compounds were compared. After using InChIKeys to retrieve the ChemOnt terms used in ClassyFire, the FDRs for the FT and TOF datasets improved to 0.0094 (19/2,027) and 0.0538 (417/7,749), respectively. Further examination of non-matching cases in the TOF dataset revealed differences in the glycosidic linkage positions in flavonoid glycosides in 231 cases (e.g., query ID PPR100244, flavonoid-3-*O*-glycosides; hit, flavonoid-7-*O*-glycosides) and leucine/isoleucine distinctions in 28 cases (e.g., query ID PPR304109, leucine and derivatives; hit, isoleucine and its derivatives) (Supplementary Data 9). Excluding cases that were nearly identical at the structural isomer level, the estimated FDR was expected to be <2%.

As strict discrimination of structural isomers by mass spectrometry alone remains challenging[5], Level 3 of the confidence scale permits tentative annotations that contain such ambiguities. Moreover, 82.8% of spectra in MassBank Human were Orbitrap mass spectrometry data (recorded as FT-MS in MassBank records). Thus, the planar structures and chemical classes of the GNPS spectral library search hits were used as pseudo-ground truths for tentative annotations in this study.

**Supplementary Note 3.** Rank aggregation using the Zipf distribution
The MS2 spectral annotation software packages output rankings of the molecular formulae or structures based on the calculated scores. We investigated the nature of these rankings and aggregated them from multiple software tools (MS-FINDER, SIRIUS, and MetFrag).
We found that the rank distributions of the correct answers followed a Zipf distribution. TOF-MS spectra were extracted from MassBank to create an evaluation dataset (12,464 spectra). Using the evaluation dataset as queries, planar structure annotations were performed using MS-FINDER to evaluate the rank positions of the correct answers. The number of hits at rank *k* was defined as $h_k$ (Supplementary Figure 2a). A log–log plot of rank k (x-axis) against frequency $h_k$ (y-axis) revealed a linear downward trend, demonstrating that the distribution of true hits obeyed Zipf's law (Supplementary Figure 2b).
The Zipf distribution is observed across various rank distributions[6], in which the frequency of occurrence at rank k is expressed as follows:

$$\mathrm{Zipf}(k,s) = k^s / \Sigma_{n=1}^{N} n^s \qquad \text{Eqn(1)}$$

The slope of the line corresponds to *s*. Similar linearity (power-law behavior) was observed for SIRIUS and MetFrag for rank distributions of molecular formulae and chemical classes and for the MassBank FT-MS dataset (Supplementary Figure 3). These results confirm that Zipf distribution can serve as a robust ranking model.
Furthermore, Zipf(*k*, *s*) can be interpreted as the likelihood at rank k is top rank (*L*). Moreover, an aggregated rank can be generated using Eqn (2), as described above.

$$L = \mathrm{Zipf}(k, s_{\mathrm{SIRIUS}}) \times \mathrm{Zipf}(l, s_{\mathrm{MS\text{-}FINDER}}) \times \mathrm{Zipf}(m, s_{\mathrm{MetFrag}}) \qquad \text{Eqn (2)}$$

The aggregated ranks were determined for the MassBank TOF-MS dataset, and Supplementary Figure 3 shows the distribution of true hits. The aggregated ranks also followed a Zipf distribution. A steeper slope compared with the individual tools indicates a better ranking performance. A similar improvement was observed in FT-MS data (Supplementary Figure 3).

**Supplementary Note 4.** Extension of annotation confidence level criteria

In this study, we performed metabolite annotation using a methodology that differed from conventional spectral matching-based workflows. Herein, we map our results to conventional confidence level criteria.

The annotation confidence level criteria proposed by Schymanski et al.[7] have been widely adopted to standardize metabolite identification in untargeted metabolomics. These criteria classify annotations into five levels, with Level 2 subdivided into two levels:

- Level 1: Confirmed structure via reference standard matching of MS, MS2, and retention time.
- Level 2a: Probable structure via exact spectral matching with library databases.
- Level 2b: Probable structure via sufficient diagnostic fragments or contextual knowledge to unequivocally pinpoint one specific isomer.
- Level 3: Tentative candidate, representing a "gray zone" of structural ambiguity (e.g., positional isomers).
- Level 4: Unequivocal molecular formula.
- Level 5: Exact mass (*m/z*) of interest without formula assignment.

Despite their widespread use, the specific evidence inventories and quantitative thresholds required for each level remain ambiguous owing to the diversity of instruments and data processing workflows across metabolomic studies. In particular, many annotations fall into Level 3, which has consequently become overly broad and heterogeneous. Several field- and platform-specific refinements and extensions have been proposed to address these limitations[7-10].

We mapped our annotations onto confidence levels (Supplementary Table 10). Among the annotation tasks performed in this study, Level 2a was assigned during the reprocessing pipeline when both a spectral match to a library spectrum (spectral entropy > 0.7) and a q-value < 0.05 were achieved by the ZBRA framework using SIRIUS, MetFrag, and MS-FINDER rankings.

For MassBank Human, we introduced four sublevels within Level 3 to explicitly differentiate the planar structure candidates, partially characterized structures, and chemical class predictions. We refer to this classification as the Tsugawa classification, named after its proposer.

- Level 3-SP (structure proposed): A single two-dimensional constitutional structure is proposed but not confirmed by a reference standard. Stereochemistry or absolute configurations may remain unresolved unless explicitly claimed.
- Level 3-SC (substructure complete): Defined molecular components, building blocks, or modifications are proposed; however, their positions, connectivities, or arrangements remain unresolved.
- Level 3-SI (substructure incomplete): A diagnostic substructure or structural motif is proposed; however, the component-level representation is not sufficiently defined.
- Level 3-CP (class proposed): A chemical or metabolite class is proposed without a stronger structure, component, or motif claim.

Supplementary Table 10 shows the evidence and thresholds used in this study.

Level 3-SP was used when a single planar structure was proposed based on the available evidence. In the reprocessing pipeline, Level 3-SP was assigned when a q-value < 0.05 was obtained by the ZBRA framework using the SIRIUS, MetFrag, and MS-FINDER rankings. Although Levels 3-SC and 3-SI were not used in this study, they would be useful for annotation based on the partial characterization of molecular structures.

Candidates were assigned to level 4 (unequivocal molecular formula) when they satisfied a q-value < 0.01 by the ZBRA framework using the SIRIUS and MS-FINDER rankings. The remaining spectra were assigned to Level 5 (exact mass of interest). The details of the annotation confidence level criteria and evidence inventories are currently being formulated and will be reported elsewhere.

**Supplementary Note 5.** Assessment of the performance of Zipf–Bayesian rank aggregation
The performance of the ZBRA method was evaluated using other approaches different from that described in the main text. The results of GNPS search and the ZBRA method were compared for 14,198 spectra curated by planar structure annotation. Among the 6,984 spectra annotated by both the GNPS search (spectral entropy > 0.7) and the ZBRA method (q-value < 0.05), 559 (8.0%) were mismatched (Supplementary Figure 4). These mismatches were derived from errors in both methods. For GNPS search, benchmarking with MassBank datasets estimated the FDRs in planar structure annotation to be 3.6% for FT-MS and 9.2% for TOF-MS (Fig. 2g and Supplementary Note 2). These results suggest that the errors derived from the ZBRA method were <5%. Similarly, the FDRs of ZBRA for the class and formula estimations were estimated to be <5% and <1%, respectively (Supplementary Figure 4).
We have found 13 unrealistic structures in 18 spectra by further examination of the structures of compounds annotated at Levels 2a or 3-SP (Supplementary Table 11). For instance, one benzyloxycarbonyl (Cbz)-protected and four *tert*-butoxycarbonyl (Boc)-protected peptides were identified by searching for compound names and SMILES strings. Protected peptides were unlikely to be present in the human samples. Moreover, alkyne-containing structures were identified by searching SMILES strings for "C#C." We also found unrealistic structures, including ynamine (C#C–NH), aryl methyl peroxide (Ar–O–O–$CH_3$), peroxyphosphate ester (P–O–O–$CH_3$), iminium ion, and free carbamic acid (R–NH–COOH). However, the natural presence of these moieties in human samples is unlikely. The removal of such compounds from large compound databases is a topic for future research. However, spectra with unrealistic structures accounted for only 0.13% (18/14,198) of planar structure annotations, which was well within the expected FDR. Warnings were issued for MassBank records with unrealistic annotations.

**Supplementary Note 6.** Structural annotation of tocopherols using the inter-dataset molecular network

As MassBank Human integrates spectra from multiple datasets, unidentified spectra that recur across many datasets represent the highest-priority cases for structure elucidation. Among the Level 4 spectra annotated only with molecular formulas, spectra similar to HSA076P0304006 were the third most widespread and were detected in 23 datasets (Supplementary Table 4). The software annotation record for HSA076P0304006 showed that the top SIRIUS hit was benzododecinium, a quaternary ammonium surfactant. This assignment is considered correct, given that the MS2 spectrum matches the fragmentation features reported in the literature, and that widespread contamination by surfactants is plausible[11]. However, annotation failed under our consensus workflow because this compound was absent from the MS-FINDER compound database.

The fifth most widespread spectra, detected in the 20 datasets, were those similar to that of HSA094P0429000. SIRIUS, MS-FINDER, and MetFrag failed to reach a consensus, ranking a vitamin E analog, cholesteryl acetate, and α-tocomonoenol at the top, respectively. Therefore, we extracted a subnetwork containing HSA094P0429000 from the molecular network shown in Extended Data Fig. 2 (Supplementary Figure 5). This subnetwork included α-tocopherol acetate (4), α-tocopherol (5), and α-tocotrienol (8) at Levels 2a and 3-SP. The fragmentation pattern of an MS2 spectrum annotated as α-tocopherol (HSA079P0431016) matched the corresponding MassBank record (MSBNK-mFam-MC20_001254). The node containing HSA094P0429000 was directly connected to α-tocopherol with a mass difference of −H2, and fragment analysis indicated dehydrogenation within the side chain. Among the possible structural isomers, we tentatively assigned the spectrum as α-tocomonoenol (6) because this isomer is the most frequently reported in humans. Because tocochromanols have numerous structural isomers, these metabolites are inherently difficult to elucidate from MS2 spectra alone. SIRIUS, MS-FINDER, and MetFrag ranked α-tocomonoenol at positions 5, 6, and 1, respectively.

We analyzed the surrounding network and tentatively assigned structures to eleven additional analogs. The metabolite series identified in this study did not match the known human catabolism of vitamin E[12]. Dietary vitamin E consists mainly of α-tocopherol (5) and α-tocotrienol (8). After absorption, the terminal side chain is ω-oxidized to carboxy intermediates in the liver, which undergo β-oxidative chain shortening and are excreted in urine. In contrast, the subnetwork in Supplementary Figure 5 contained hydroxylated (13, 15), carboxylated (12, 14), and β-oxidized (18) forms derived not from α-tocopherol or α-tocotrienol but from α-tocomonoenol and α-tocodienol. Moreover, α-tocopherol and α-tocomonoenol were detected across diverse sample types, whereas all other analogs were detected exclusively in fecal samples. This skewed distribution suggests that these analogs arise from gut bacterial hydrogenation or dehydrogenation of dietary α-tocopherol and α-tocotrienol. Thus, the interplay between dietary input, host metabolism, and gut microbial transformation appears to generate metabolite diversity, even for vitamin E.

Achieving such detailed annotations requires detailed spectral comparisons within molecular networks. No single dataset contained all the metabolites in this network; their detection and interpretation became possible only through an inter-dataset molecular network integrating numerous datasets. Thus, MassBank Human will guide the exploration of uncharted chemical spaces in the human metabolome.

**References**


65 Nakatani, K., Izumi, Y., Takahashi, M. & Bamba, T. Unified-hydrophilic-interaction/Anion-exchange liquid chromatography mass spectrometry (Unified-HILIC/AEX/MS): A single-run method for comprehensive and simultaneous analysis of polar metabolome. Anal. Chem. 94, 16877-16886 (2022).

66 Zhang, P. et al. Revisiting fragmentation reactions of protonated alpha-amino acids by high-resolution electrospray ionization tandem mass spectrometry with collision-induced dissociation. Sci Rep 9, 6453 (2019).

67 Kind, T. & Fiehn, O. Advances in structure elucidation of small molecules using mass spectrometry. Bioanalytical reviews 2, 23-60 (2010).

68 Koelmel, J. P. et al. An actionable annotation scoring framework for gas chromatography-high-resolution mass spectrometry. Exposome 2, osac007 (2022).

69 Charbonnet, J. A. et al. Communicating confidence of per- and polyfluoroalkyl substance identification via high-resolution mass spectrometry. Environ Sci Technol Lett 9, 473-481 (2022).

70 Celma, A. et al. Ion mobility separation coupled to high-resolution mass spectrometry in environmental analysis – Current state and future potential. Trends in Environmental Analytical Chemistry 43, e00239 (2024).

71 Watrobska-Swietlikowska, D. Distribution of benzalkonium chloride into the aqueous phases of submicron dispersed systems: emulsions, aqueous lecithin dispersion and nanospheres. AAPS PharmSciTech 21, 7 (2019).

72 Schmolz, L., Birringer, M., Lorkowski, S. & Wallert, M. Complexity of vitamin E metabolism. World J Biol Chem 7, 14-43 (2016).

### Supplementary Figures

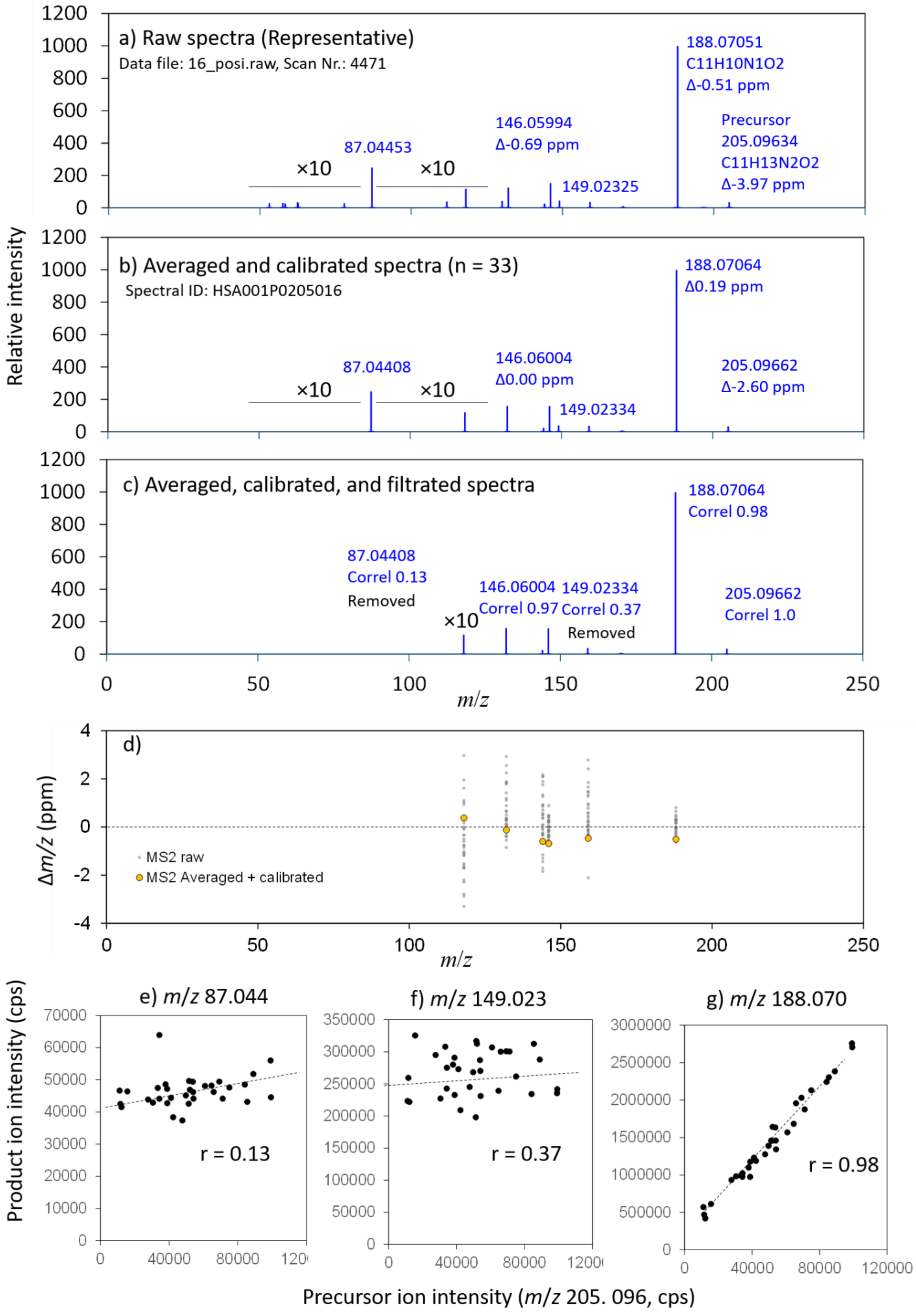

**Supplementary Figure 1.** An example of preprocessing. This figure details how the three-step preprocessing (averaging, recalibration, and filtering) transforms raw, noisy DDA data into a high-quality spectral record, illustrating the generation of HSA001P0205016. Spectra were obtained from tryptophan.
(a–c) MS2 spectra at each processing step: a representative raw spectrum (a), the averaged and recalibrated spectrum (b), and the final filtered spectrum (c). Preprocessing retained reproducible fragment ions while removing noise (e.g., $m/z$ 87.044) and contaminants.
(d) Improvement in mass accuracy. Mass errors ($\Delta m/z$) are plotted across the $m/z$ range. Averaging and recalibration (orange) substantially reduced the dispersion compared with that in the raw data (gray), achieving sub-ppm accuracy.
(e–g) Intensity correlations used for filtering. Scatter plots of precursor and product ion intensities. A true fragment (g) shows a high correlation ($r = 0.98$), whereas interference signals (e, f) show low correlations ($r < 0.40$) and were subsequently filtered.

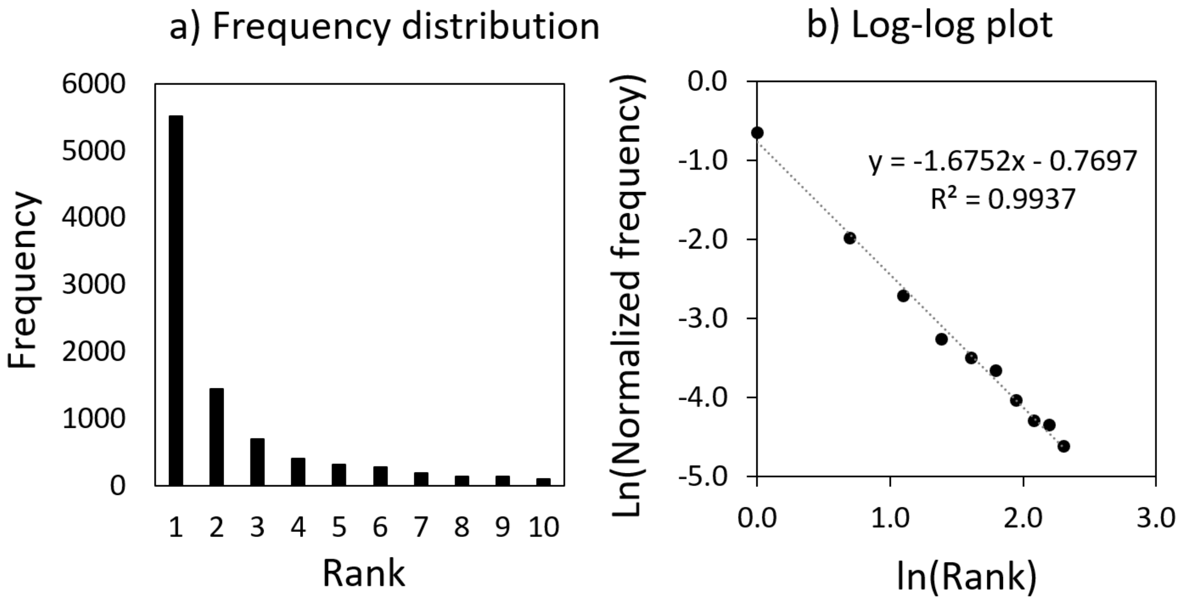


**Supplementary Figure 2.** Frequency distribution of correct hits in the planar structure ranking by MS-FINDER. The MassBank TOF-MS dataset (12,464 spectra) was used as query spectra. (a) Frequency distribution of correct hits. (b) Log–log plot. Ranks 1–10 were used for the analysis.

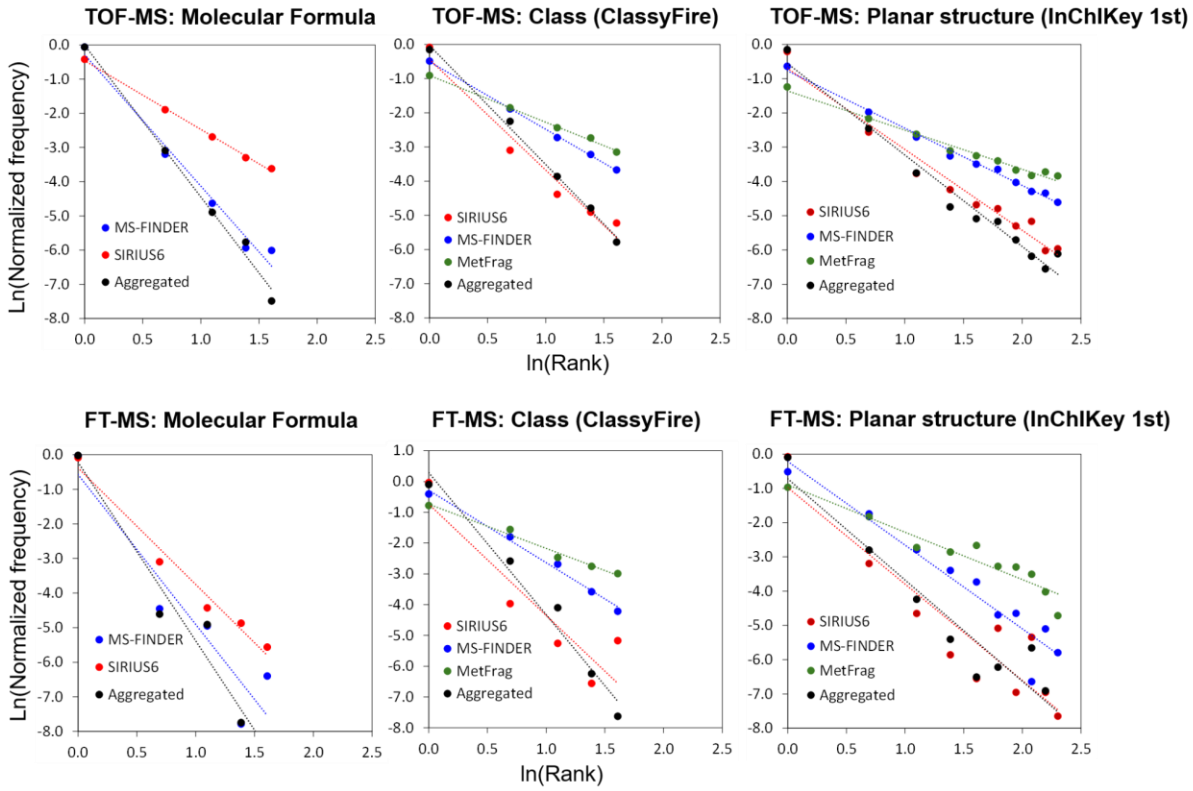


**Supplementary Figure 3**. Frequency distribution of correct hits in the molecular formula, class, and planar structure rankings by MS-FINDER, SIRIUS, MetFrag, and their aggregates. The MassBank TOF-MS (12,464 spectra) and FT-MS (3,234 spectra) datasets were used as query spectra. Ranks 1–5 were used in the molecular formula and class analyses to avoid infrequent data, whereas ranks 1–10 were used in the planar structure analysis.

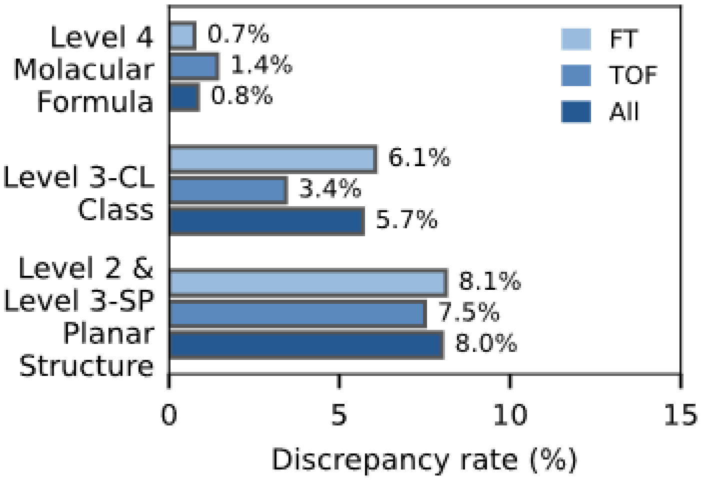


**Supplementary Figure 4.** Evaluation of the FDR through comparison with GNPS library search results.

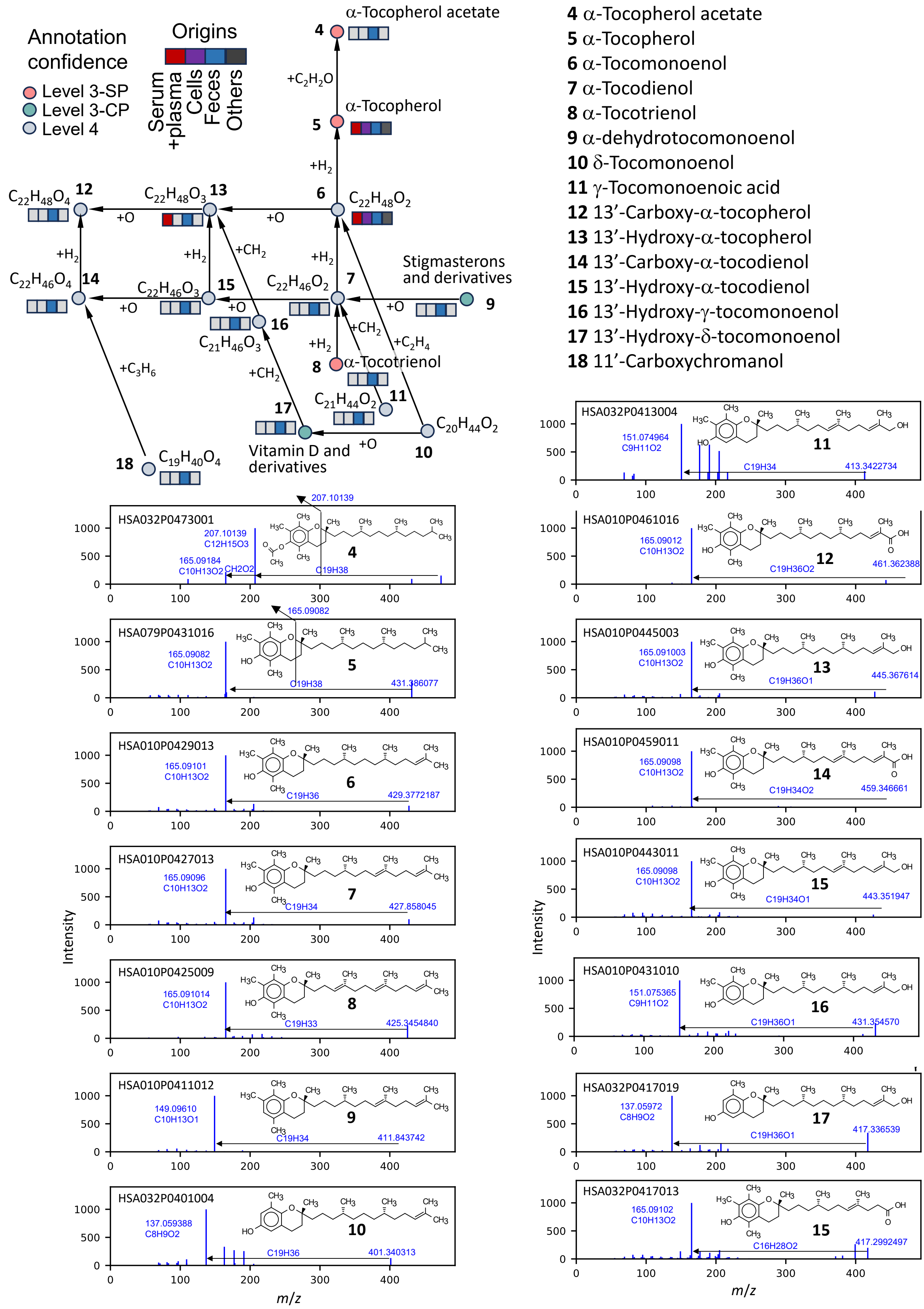


**Supplementary Figure 5.** Structural elucidation of α-tocopherol derivatives (vitamin E) using the inter-dataset molecular network. The molecular network and representative MS2

spectra are shown together with the deduced structures. To examine these spectra, the MassBank Human database provides an SMB_Measured ID search service (https://human.massbank.jp/SMB_ID_Search_page), through which details of the spectral data can be retrieved by record ID. Please add the header “MSBNK-SMB_Measured-” to all IDs such as MSBNK-SMB_Measured-HSA032P0473001.

**4**, α-tocopherol acetate, HSA032P0473001; **5**, α-tocopherol, HSA079P0431016; **6**, α-tocomonoenol, HSA010P0429013; **7**, α-tocodienol, HSA010P0427013; **8**, α-tocotrienol, HSA010P0425009; **9**, α-dehydrotocomonoenol, HSA010P0411012; **10**, δ-tocomonoenol, HSA032P0401004; **11**, γ-tocomonoenoic acid, HSA032P0413004; **12**, 13′-carboxy-α-tocopherol, HSA010P0461016; **13**, 13′-hydroxy-α-tocopherol, HSA010P0445003; **14**, 13′-carboxy-α-tocodienol, HSA010P0459011; **15**, 13′-hydroxy-α-tocodienol, HSA010P0443011; **16**, 13′-hydroxy-γ-tocomonoenol, HSA010P0431010; **17**, 13′-hydroxy-δ-tocomonoenol, HSA032P0417019; **18**, 11′-carboxychromanol, HSA032P0417013
